# DFFB suppresses interferon to enable cancer persister cell regrowth

**DOI:** 10.1101/2025.08.15.670603

**Authors:** August F. Williams, David A. G. Gervasio, Claire E. Turkal, Anna E. Stuhlfire, Michael X. Wang, Brandon E. Mauch, Rhea Plawat, Ariel H. Nguyen, Michelle H. Paw, Mehrshad Hairani, Cooper P. Lathrop, Sophie H. Harris, Jennifer L. Page, Matthew J. Hangauer

**Affiliations:** Department of Dermatology, School of Medicine, University of California San Diego; Moores Cancer Center, University of California San Diego; Altman Clinical and Translational Research Institute, University of California San Diego; Stem Cell Core, Salk Institute for Biological Studies, La Jolla, CA

## Abstract

Oncogene targeted cancer therapies can provide deep responses but frequently suffer from acquired resistance.^1^ Therapeutic approaches to treat tumours which have acquired drug resistance are complicated by continual tumour evolution and multiple co-occurring resistance mechanisms.^2,3^ Rather than treating resistance after it emerges, it may possible to prevent it by inhibiting the adaptive processes which initiate resistance but these are poorly understood.^4^ Here we report that residual cancer persister cells that survive oncogene targeted therapy are growth arrested by drug stress-induced intrinsic Type I interferon (IFN) signaling. To escape growth arrest, persister cells leverage apoptotic machinery to transcriptionally suppress interferon-stimulated genes (ISGs). Mechanistically, persister cells sublethally engage apoptotic caspases to activate DNA endonuclease DNA Fragmentation Factor B (DFFB, also known as Caspase-Activated DNase (CAD)) which induces DNA damage, mutagenesis, and stress response factor Activating Transcription Factor 3 (ATF3). ATF3 limits Activator Protein-1 (AP1)-mediated ISG expression sufficiently to allow persister cell regrowth. Persister cells deficient in DFFB or ATF3 exhibit high ISG expression and are consequently unable to regrow. Therefore, sublethal apoptotic stress paradoxically promotes regrowth of residual cancer cells that survive drug treatment.

---

To discover adaptive mechanisms that enable cancer cells to acquire drug resistance, we utilized models in which treatment with oncogene targeted therapies results in death for the majority of cells and effectively purifies a residual population of primarily quiescent cancer persister cells (Extended Data Fig. 1a-d). A minority of persister cells regrow into drug-tolerant expanded persister (DTEP) cell colonies during the first three months of treatment, modeling the earliest stage of tumour recurrence (Fig. 1a and Extended Data Fig. 1a-d). Though DTEP colonies have been reported to eventually acquire resistance mutations and become irreversibly drug resistant after six or more months of treatment,^5,6^ it is not known whether mutations contribute to initial DTEP colony formation.

**Fig. 1.**
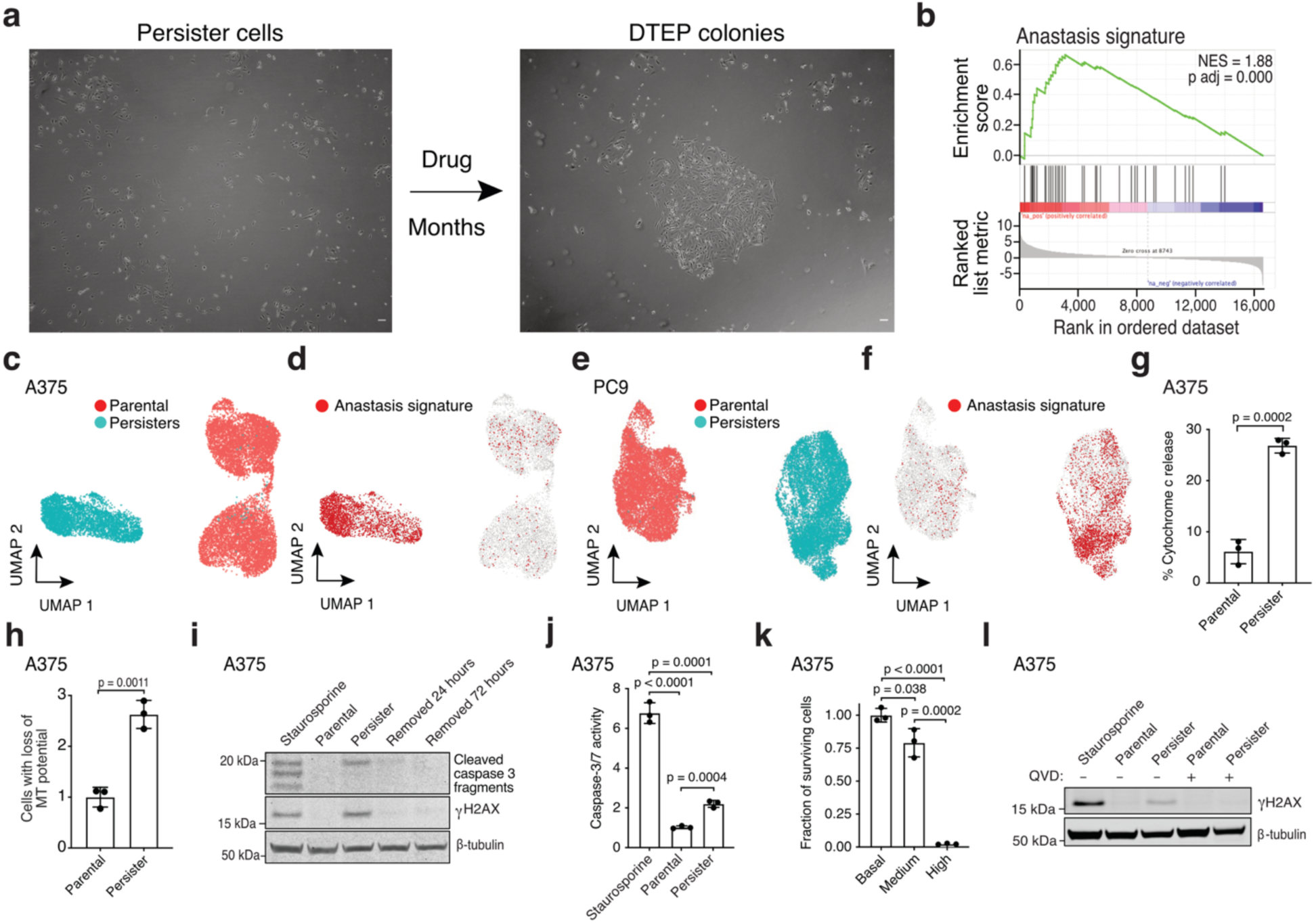
Drug-tolerant persister cells undergo chronic sublethal apoptotic stress-induced DNA damage. **a**, Microscopy of A375 persister and DTEP cells treated with 250 nM dabrafenib and 25 nM trametinib. **b**, Bulk RNAseq gene set enrichment analysis of the anastasis signature in A375 persister cells. Normalized enrichment score, NES. P value calculated with a permutation test with family-wise error rate correction. **c-f**, ScRNAseq UMAPs of A375 (**c**,**d**) and PC9 (**e**,**f**) parental cells and persister cells. UMAP, uniform manifold approximation and projection. **d**,**f**, Cells with enriched anastasis gene set are highlighted in red. **g**, A375 parental and persister cells assessed for mitochondrial release of cytosolic cytochrome c. **h**, A375 parental and persister cells assessed for loss of mitochondrial (MT) potential using JC-1 indicator dye by flow cytometry. **i**, A375 persister cells assessed for cleaved caspase 3 and γH2AX either on drug or following 24 or 72 hours of drug removal. **j**, Flow cytometry quantification of caspase 3/7 activity reporter geometric means. **k**, A375 persister cells sorted for basal, medium (sublethal), and high (lethal) caspase 3/7 activity, replated in media without drug, and assessed for cell viability the following day. **l**, Parental and persister cells assessed for γH2AX with or without co-treatment with 10 μM caspase inhibitor QVD. **i**,**j**,**l**, Parental cells treated for 24 hours with 250 nM **(i**,**l)** or 4 hours with 1 μM staurosporine **(j)** were used as a positive control. **g**,**h**,**j**,**k** *n* = 3 biological replicates; mean ± s.d. is shown; P values calculated with two-tailed Student’s t-test unless stated otherwise.

To gain insight into the transition from persister to DTEP cells (Fig. 1a), we performed RNA sequencing on targeted therapy-treated persister cells. Beyond previously described persister cell signatures,^5,7–11^ we observed an enrichment in an “anastasis” signature (Fig. 1b-f and Supplementary Tables 1-3). Anastasis is the phenomenon wherein cells recover from brief periods of apoptotic stress and this can result in DNA damage, mutagenesis, and transformation.^12^ Similarly, “flatliner” cells which survive transient mitochondrial outer membrane permeabilization (MOMP), a key initiating step toward apoptosis, were recently shown to have increased potential to form persister cells and also to metastasize.^13^ Melanoma cells which undergo “failed apoptosis” also gain migration and invasion properties.^14^ Therefore, brief encounters with apoptotic stress can have pro-tumour effects. Whether cancer cells can also tolerate weeks or months of continuous apoptotic stress such as in the context of acquired drug resistance is unknown.

## Persister cells undergo chronic sublethal apoptotic signaling

We found that persister cells exhibit partial mitochondrial cytochrome c release (Fig. 1g and Supplementary Fig. 2) and loss of mitochondrial potential (Fig. 1h and Supplementary Fig. 3) indicative of sublethal MOMP. Persister cells also have partially cleaved apoptotic executioner caspase 3 which is at a level below cells treated with a lethal exposure of apoptosis inducer staurosporine, consistent with sublethal caspase activation (Fig. 1i, Extended Data Fig. 2a, and Supplementary Fig. 4). Furthermore, using a fluorescent caspase 3/7 activity sensor, we observed an increase in caspase activity in live persister cells (Fig. 1j and Supplementary Fig. 5) and sorted persister cells with significantly elevated caspase 3/7 activity levels were capable of regrowth upon drug removal, while cells with very high caspase activity were not, demonstrating that persister cells remain viable with apoptotic caspase activity (Fig. 1k and Extended Data Fig. 2b). This is consistent with a recent study showing that cells can recover from intermediate but not high apoptotic caspase 3 activity.^15^ Also, though cancer cell lines can have spontaneous low level apoptotic signaling,^16^ we found that persister cell apoptotic signaling is drug-induced because apoptotic signaling markers are absent prior to treatment, induced early during drug exposure, maintained throughout months of treatment, and dissipate within 24 hours of drug removal (Fig. 1i and Extended Data Fig. 2c,d). Therefore, persister cells experience drug stress-induced apoptotic signaling throughout drug exposure yet remain alive.

## DFFB induces DNA damage in persister cells

In response to a variety of acute stresses, apoptotic caspases can promote DNA damage by activating apoptotic DNA endonucleases.^12,17–19^ We tested whether apoptotic caspases also induce DNA damage in persister cells and found that the pan-caspase inhibitor Quinoline-Val-Asp-Difluorophenoxymethylketone (QVD) blocked persister cell DNA damage across multiple tumour types treated with targeted therapies (Fig. 1l, Extended Data Fig. 2e-h and Supplementary Fig. 6). Similarly, caspase 9 knockout (KO) persister cells, which do not undergo drug-induced caspase 3 cleavage, fail to acquire DNA damage (Extended Data Fig. 2i). We also found that persister cells sorted for elevated caspase 3/7 activity have increased DNA damage compared to persister cells without elevated caspase activity (Extended Data Fig. 2j). Therefore, targeted therapy drugs induce apoptotic caspase-dependent DNA damage in persister cells.

During apoptosis, DNA endonuclease DFFB is activated by caspase 3-mediated proteolytic cleavage of DFFB chaperone and inhibitor protein DNA Fragmentation Factor A (DFFA, also known as ICAD or DFF45), enabling DFFB to form a nuclease active dimer.^20^ Activated DFFB induces DNA breaks promoting extensive fragmentation of chromosomal DNA during apoptotic cell death,^21–23^ but in response to transient sublethal apoptotic stress DFFB can also induce acute DNA damage.^17–19,24^ We explored whether the chronic apoptotic stress we observed in persister cells results in DFFB-mediated persistent DNA damage. Consistent with activation of DFFB, we observed cleavage of DFFA in persister cells (Fig. 2a) which is dependent on caspases (Extended Data Fig. 2i,k). DFFA cleavage was also observed in DTEP cells indicating continuous DFFB activation throughout DTEP colony formation (Extended Data Fig. 2k). To test whether DFFB functionally contributes to DNA damage, we generated DFFB loss of function (LOF) models including CRISPR-mediated DFFB KO cells (Extended Data Fig. 3a-g) and cells with ectopically expressed noncleavable mutant DFFA which is resistant to caspase-mediated cleavage and blocks endogenous DFFB activity.^25^ We found that drug-induced DNA damage was absent in all DFFB LOF persister cell models (Fig. 2b-e and Extended Data Fig. 3h) and re-expression of WT DFFB, but not nuclease activity deficient mutant DFFB,^26^ restored drug-induced DNA damage in DFFB KO persister cells (Fig. 2f). Therefore, DFFB endonuclease activity is the primary source of drug-stress induced DNA damage in targeted therapy treated persister cells.

**Fig. 2.**
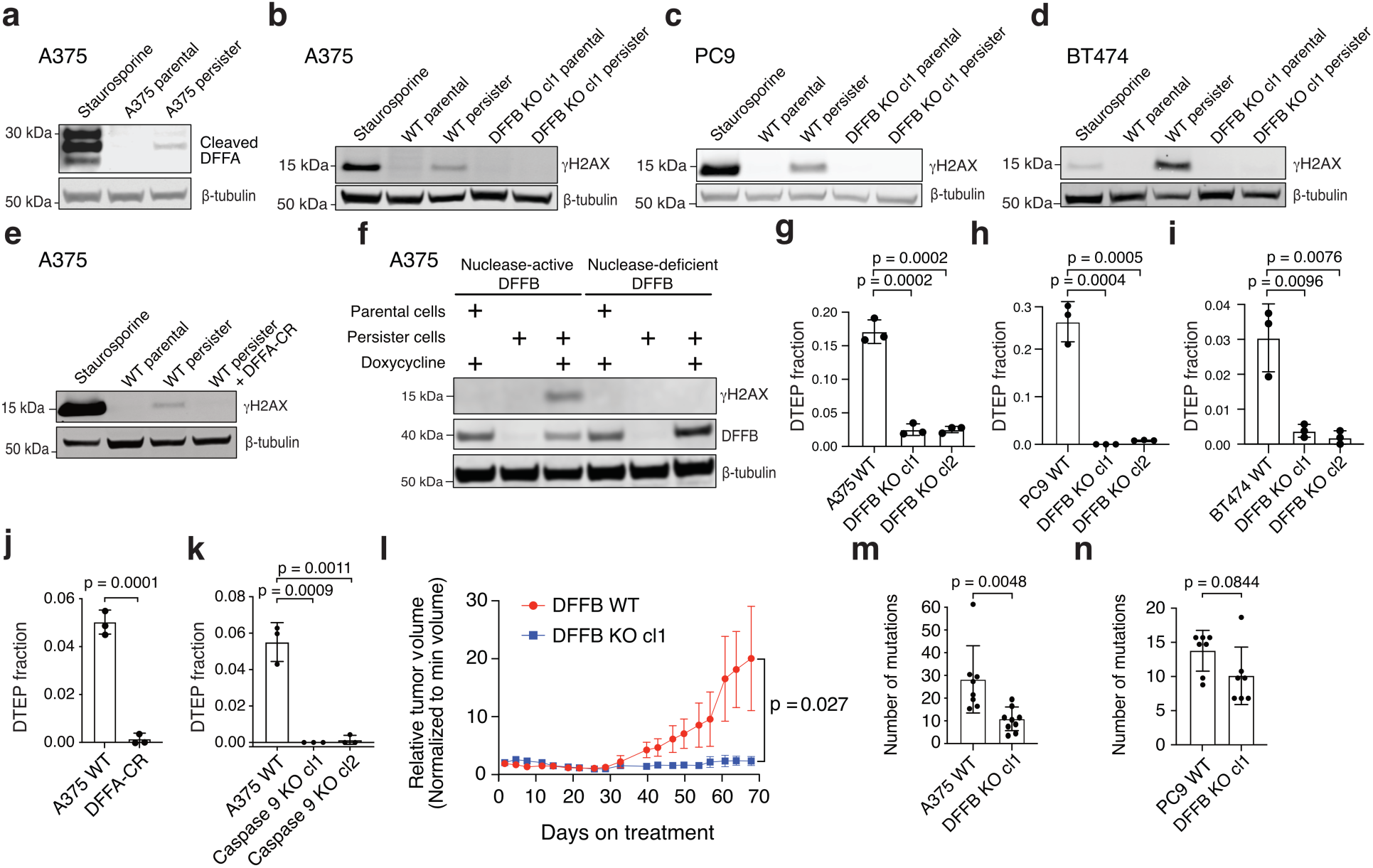
Apoptotic DNase DFFB mediates persister cell DNA damage, mutagenesis and regrowth. **a**, A375 persister cells exhibit cleaved DFFA indicating DFFB activation. **b-e**, Persister cell DNA damage is absent in A375 (**b**,**e**), PC9 (**c**), and BT474 (**d**) DFFB loss of function (LOF) persister cells. **a**-**e**, 250 nM (**a**,**b**,**e**), 500 nM (**c**), or 750 nM (**d**) staurosporine treatment for 24 hours was used as a positive control. cl, clone. DFFA-CR, ectopically expressed DFFA cleavage resistant variant. **f**, Doxycycline-inducible re-expression of nuclease active DFFB, but not nuclease dead DFFB, in DFFB KO A375 persister cells restores drug-induced DNA damage. **g-k**, Measurement of fraction of cells which regrow into DTEP colonies during treatment. *n* = 3 biological replicates; mean ± s.d.; two-tailed Student’s t-test. **g**,**j**,**k**, A375 melanoma cells treated with dabrafenib and trametinib. **h**, PC9 lung cancer cells treated with osimertinib. **i**, BT474 breast cancer cells treated with lapatinib. **l**, Tumour volume of A375 xenograft tumours in mice treated with dabrafenib and trametinib. Data are normalized to the minimum volume of each tumour during treatment; *n* = 3 biological replicates; mean ± s.d.; two-tailed Student’s t-test. **m**,**n**, Whole exome sequencing analysis of mutations acquired by A375 cells during 7 weeks of dabrafenib and trametinib treatment (**m**) and PC9 cells during 10 weeks of 2.5 μM erlotinib treatment (**n**). **m**, *n* = 8 WT biological replicates and 9 DFFB KO biological replicates. **n**, *n* = 7 WT and 7 DFFB KO biological replicates. **m**,**n**, Mean ± s.d.; two-tailed Student’s t-test.

## DFFB is required for persister cell regrowth and tumour relapse

Given prior observations that DFFB activation can promote transformation,^18,27^ we explored whether chronic DFFB-induced DNA damage may promote persister cell escape from growth arrest to enable regrowth into DTEP colonies. Indeed, we found that DFFB LOF persister cells were significantly hindered in their ability to form DTEP colonies across all tested tumour types and treatments (Fig. 2g-k and Extended Data Fig. 3i). We then tested whether DFFB is required for acquired resistance in vivo. Mice bearing A375 DFFB WT or KO melanoma tumours, which formed similarly (Extended Data Fig. 4a), were treated with dabrafenib and trametinib. Both DFFB WT and KO tumours initially responded to therapy and shrunk to a similar minimal residual volume (Extended Data Fig. 4b). However, while DFFB WT tumours regrew during further treatment, DFFB KO tumours remained in a regressed state and failed to regrow (Fig. 2l). These observations are independent of tumour cell viability, initial drug response or persister cell formation which were each found to be unaffected by DFFB deficiency (Extended Data Fig. 4). Therefore, DFFB is specifically required for persister cells and residual tumours to regrow during drug treatment.

## DFFB induces mutagenesis in persister cells

We next explored the mechanism by which DFFB enables persister cell regrowth. One possible mechanism is by inducing mutagenesis and acquisition of resistance mutations because DFFB has been reported to promote mutagenesis during acute apoptotic stress.^17,27,28^ To test this possibility, we performed whole exome sequencing (WES) of DFFB WT and KO A375 cells subjected to 7 weeks of dabrafenib and trametinib treatment which revealed a variety of acquired mutations, supporting recent reports of persister cell stress-induced mutagenesis (Supplementary Table 4).^7,29^ Consistent with DFFB-mediated mutagenesis, we found that A375 DFFB KO cells had fewer than half as many acquired mutations as WT cells (Fig. 2m). Analysis of PC9 cells subjected to 10 weeks of erlotinib treatment similarly revealed that PC9 DFFB KO cells acquired fewer mutations than WT cells (Fig. 2n and Supplementary Table 5). However, the magnitude of the contribution of DFFB to acquired mutations was less in PC9 potentially because PC9 persister cells also undergo significant APOBEC3-mediated mutagenesis^29^ and lack downregulation of DNA repair genes or upregulation of error prone DNA polymerases, a previously reported feature of stress-induced mutagenesis^7^ (Extended Data Fig. 5a-e). These data suggest DFFB is one of multiple sources of mutagenesis in persister cells. However, there were no known resistance conferring mutations found among the acquired mutations (Supplementary Tables 4 and 5). We therefore searched for an alternative nonmutational mechanism by which DFFB promotes persister cell regrowth.

## DFFB suppresses interferon signaling to enable persister cell regrowth

RNAseq analysis of WT persister and DTEP cells revealed significantly enriched signatures composed of IFN-stimulated genes (ISGs) in both populations compared to parental cells, though ISGs were decreased in cycling DTEP cells (Fig. 3a, Extended Data Fig. 6a-d, and Supplementary Table 2). Tumour cell-intrinsic IFN signaling can contribute to growth arrest^30^ and suppress disseminated cancer cell growth,^31^ and therefore could limit persister cell regrowth. One potential source of persister cell intrinsic IFN is MOMP which, in apoptotic cells, exposes mitochondrial nucleic acids to cytoplasmic pattern recognition receptors that induce type I IFN.^32^ We found that persister cells, which exhibit MOMP (Fig. 1g,h), also have elevated exposure of mitochondrial nucleic acids to cytosol and upregulated IFNβ expression (Extended Data Fig. 6e,f). We therefore hypothesized that persister cell regrowth is limited by intrinsic IFN signaling. While a minority of WT persister cells regrow into DTEP colonies, most WT persister cells do not (Fig. 2g-k) and we found that co-treatment of persister cells with JAK inhibitor ruxolitinib to block IFN signaling increased the proportion of persister cells which regrew into DTEP colonies (Extended Data Fig. 6g,h). Furthermore, DTEP colonies are predominantly constituted by quiescent cells with elevated ISGs, indicating that ISGs remain growth suppressive throughout treatment (Extended Data Fig. 6i-l).

**Fig. 3.**
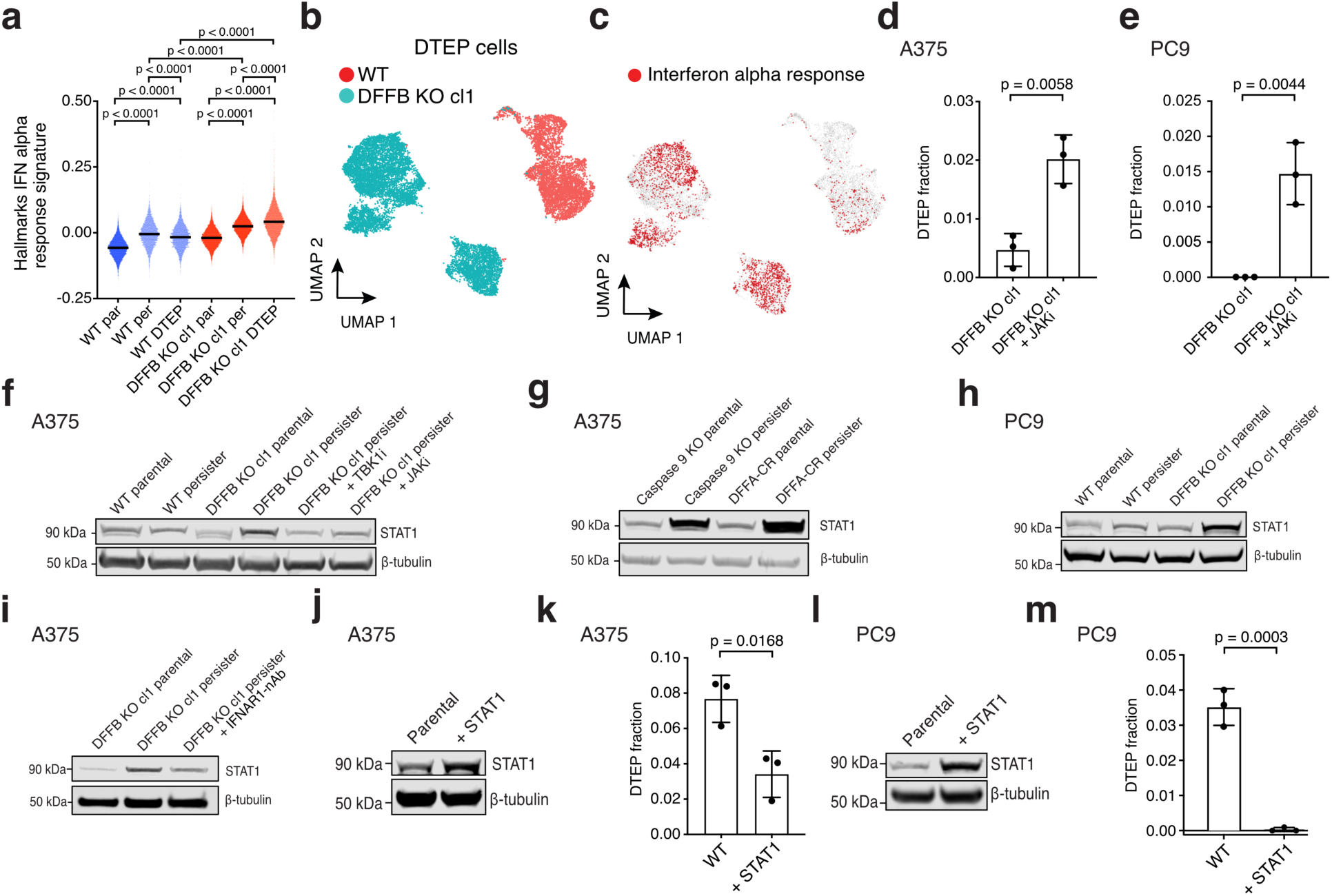
DFFB suppresses interferon signaling to allow persister cell regrowth. **a**, ScRNAseq Hallmarks IFN alpha response gene set signature scores in A375 DFFB WT and KO parental cells and drug treated cells at the persister and DTEP timepoints. P values calculated with Mann-Whitney test. cl, clone. **b-c**, ScRNAseq UMAP of similar numbers of A375 WT and DFFB KO cells at 9 weeks of drug treatment (**b**) with enriched Hallmarks interferon alpha response gene set highlighted in red (**c**). **d**,**e**, Measurement of fraction of cells which regrew into DTEP colonies during treatment. **d**, A375 DFFB KO cells treated with 250 nM dabrafenib and 25 nM trametinib for 2 weeks to derive persister cells, then 1 μM JAK inhibitor ruxolitinib was added and all three drugs were maintained for 5 more weeks. **e**, PC9 DFFB KO cells treated with 300 nM osimertinib and 5 μM ruxolitinib for 5 weeks. **f**, Total STAT1 levels in A375 persister cells co-treated for 14 days with 1 μM TBK1 inhibitor MRT67307 or 1 μM JAK inhibitor**. g**, A375 caspase 9-depleted (pooled KO) and cleavage resistant DFFA (DFFA-CR)-expressing parental and persister cells analyzed for STAT1. **h**, PC9 WT and DFFB KO persister cells treated with 500 nM osimertinib analyzed for STAT1. **i**, A375 DFFB KO cells treated for 7 days with 250 nM dabrafenib and 25 nM trametinib with and without 1.5 μM of interferon alpha receptor 1 neutralizing antibody (IFNAR1-nAb) and analyzed for total STAT1 expression. **j**,**k**, A375 WT and STAT1 overexpressing cells treated with 250 nM dabrafenib and 25 nM trametinib and analyzed for DTEP colony formation. **l**,**m**, PC9 WT and STAT1 overexpressing cells treated with 300 nM osimertinib and analyzed for DTEP colony formation. **d**,**e**,**k**,**m**, *n* = 3 biological replicates; mean ± s.d. is shown; P values calculated with two-tailed Student’s t-test.

Interestingly, we found that drug treated DFFB KO cells exhibit strongly elevated IFN transcriptional signatures above corresponding WT cells at all timepoints (Fig. 3a-c; Extended Data Fig. 6m-o; and Supplementary Tables 6-9). We also observed stronger rescue of DTEP colony formation in DFFB KO than WT persister cells upon treatment with JAK inhibitor (Fig. 3d,e, Extended Data Fig. 6g,h). Furthermore, STAT1, a central IFN pathway transcription factor which drives prolonged expression of ISGs and is itself an ISG,^33,34^ is upregulated throughout drug treatment in DFFB LOF cells compared to WT cells in a type I IFN signaling-dependent manner (Fig. 3f-i and Extended Data Fig. 6p). Elevated STAT1 is sufficient to block persister cell regrowth because STAT1 ectopic overexpression suppresses WT DTEP formation (Fig. 3j-m). In contrast, elevated STAT1 has minimal effect on parental cell proliferation suggesting that IFN signaling in the absence of drug stress is insufficient to block growth (Extended Data Fig. 6q,r). These observations demonstrate that during drug treatment, DFFB suppresses IFN signaling sufficiently to allow regrowth of a minority of persister cells. Consequently, DFFB-deficient persister cells experience highly elevated IFN signaling which prevents regrowth.

We then explored how DFFB regulates IFN signaling. Though high levels of DFFB-induced DNA damage were recently reported to promote transient STING-mediated type I IFN production during direct ectopic DFFB activation or acute viral infection,^35^ persister cells have lower levels of drug stress-mediated DFFB-induced DNA damage and this is insufficient to drive DFFB-dependent IFN induction (Extended Data Fig. 6f,s). Type I IFN production is instead regulated independent of DFFB in persister cells by apoptotic caspases as shown by increased IFNβ expression upon QVD treatment (Extended Data Fig. 6f), consistent with prior reports in other contexts.^36,37^ Given that ISGs are strongly elevated in DFFB KO persister cells, but expression of type I IFN is not, we hypothesized that DFFB transcriptionally regulates ISGs downstream of IFN signaling. Though DFFB is a nuclease which has some preference for inducing DNA breaks in specific genomic regions^38^ and DFFB-induced DNA damage can affect proximal gene expression,^39^ we considered it unlikely that DFFB induces coordinated suppressive DNA breaks at the numerous ISG loci DFFB regulates in persister cells. Therefore, we instead searched for an ISG suppressing factor which is activated by DFFB.

## DFFB activates stress response factor ATF3

We found that stress response factor ATF3 is specifically induced in drug treated WT cells at both the persister and DTEP timepoints (Fig. 4a and Extended Data Fig. 7a-c). ATF3 has previously been shown to either negatively or positively regulate expression of ISGs including STAT1 in other contexts.^40–43^ Therefore, we considered ATF3 to be a candidate mediator of DFFB-induced ISG suppression. We first tested how ATF3 expression is induced in persister cells. ATF3 can be induced by a variety of stresses and signals including DNA damage,^44^ IFN signaling,^45^ and the integrated stress response (ISR).^46^ ATF3 is induced in WT persister cells which have DNA damage and is not induced in DFFB KO persister cells which lack DNA damage (Extended Data Fig. 7d). Furthermore, ATF3 expression is rescued in DFFB KO persister cells by treatment with topoisomerase inhibitor etoposide which induces DNA breaks independent of DFFB^47^ thereby demonstrating that DNA damage drives ATF3 induction in persister cells (Fig. 4b). We also found that JAK inhibitor co-treatment reduced ATF3 levels in persister cells indicating IFN signaling also promotes ATF3 induction (Fig. 4c and Extended Data Fig. 7e).

**Fig. 4.**
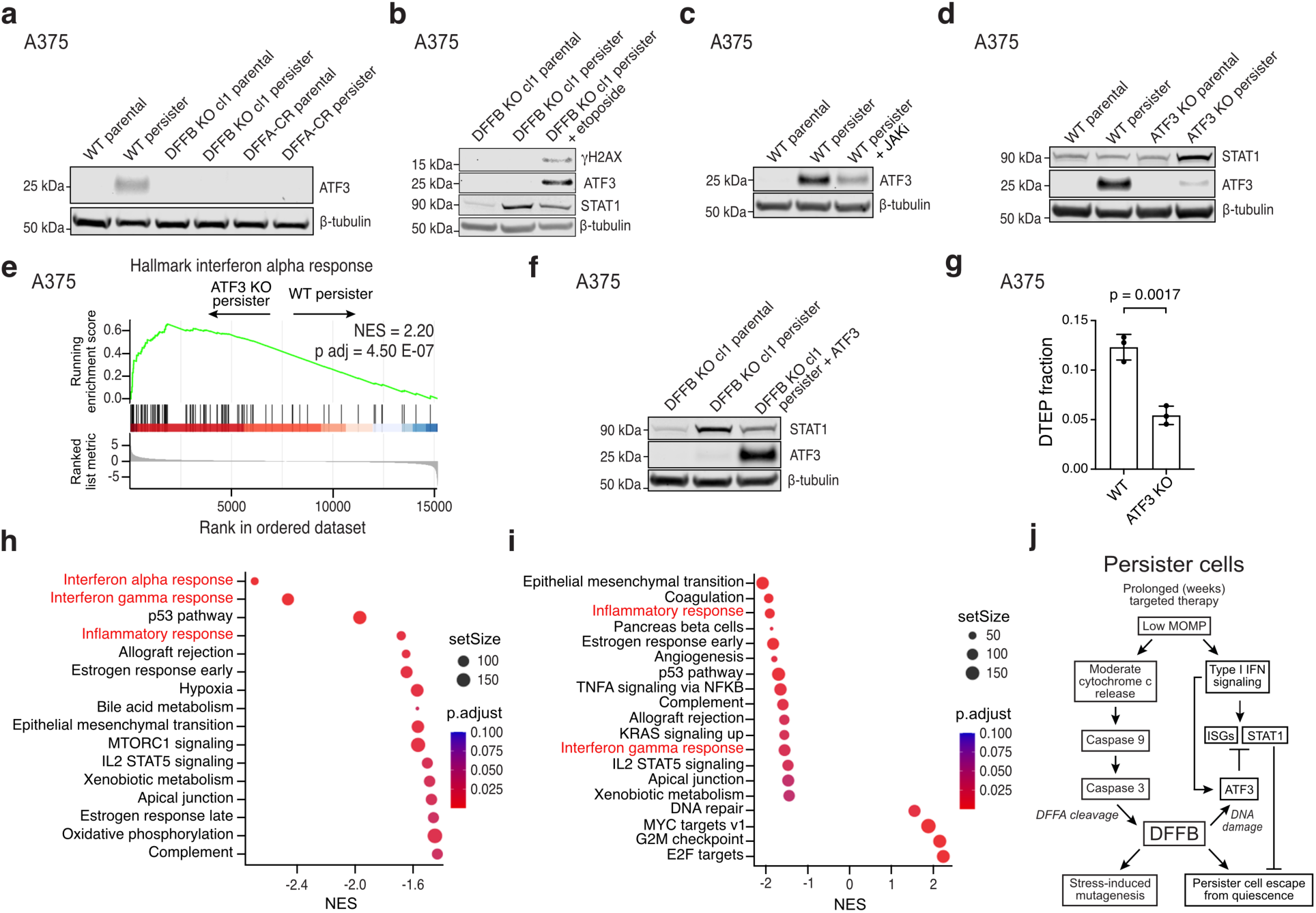
DFFB induces ATF3 to suppress ISGs and enable persister cell regrowth. **a**, ATF3 protein levels in A375 WT, DFFB KO, and DFFA-CR parental and persister cells. cl, clone. **b**, A375 DFFB KO persister cells treated with or without 100 μM etoposide for 48 hours and analyzed for DNA damage, ATF3 and STAT1. **c**, A375 WT persister cells co-treated with 1 μM JAKi and analyzed for ATF3 expression. **d**, Total STAT1 and ATF3 levels in A375 cells with CRISPR-mediated ATF3 depletion (pooled KO). **e**, Bulk RNAseq gene set enrichment analysis of Hallmarks interferon alpha response gene set using differentially expressed genes between A375 ATF3 KO and WT persister cells. Normalized enrichment score, NES. **f**, Total STAT1 and ATF3 protein levels in A375 DFFB KO persister cells with ectopically expressed ATF3. **g**, DTEP colony formation in A375 WT and ATF3-depleted (pooled KO) cells. *n* = 3 biological replicates; mean ± s.d.; two-tailed Student’s t-test. **h**-**i**, Bulk RNAseq gene set enrichment analysis of A375 ATF3 KO (**h**) and DFFB KO (**i**) persister cells with versus without co-treatment with 20 μM AP1 inhibitor T-5224. Interferon-related gene sets are colored red. *n* = 3 biological replicates. **j**, Summary diagram.

The ISR is activated by a variety of stresses resulting in phosphorylation of eIF2α and induction of ATF4 which drives expression of target genes including ATF3.^48^ Recently, the ISR was shown to also be induced via HRI activation as a result of MOMP and cytochrome c release in cancer cells which recover from transient treatment with cytotoxic concentrations of BH3 mimetics, resulting in cells with persister cell features and metastatic potential.^13^ We also observed that transient treatment with BH3 mimetics induces the ISR and induction of ATF3 in recovered A375 and PC9 cells (Extended Data Fig. 7f,g). However, we failed to detect phosphorylated eIF2α or increased ATF4 at any timepoint during targeted therapy treatment including in persister cells (Extended Data Fig. 7d,f,g). An explanation for this distinction is that cytotoxic BH3 mimetic-treated cells exhibit much higher mitochondrial cytochrome c release compared to targeted therapy treated persister cells, suggesting that a minimum threshold of cytoplasmic cytochrome c is required to activate the ISR (Extended Data Fig. 7h-k and Supplementary Fig. 7). Furthermore, while ATF3 induction is dependent on ATF4 during the ISR, we found that in persister cells ATF3 is induced independently of ATF4 because ATF4 KO persister cells robustly induce ATF3 (Extended Data Fig. 7l,m). Therefore, DFFB-dependent ATF3 induction in persister cells is mediated by DNA damage and IFN signaling, not the ISR.

## ATF3 suppresses ISG expression to promote persister cell regrowth

We next tested whether ATF3 functionally suppresses ISGs in persister cells to allow regrowth and found that CRISPR-mediated ATF3 depletion resulted in increased STAT1 and elevated IFN-related gene sets (Fig. 4d,e and Supplementary Tables 10 and 11). Furthermore, in DFFB KO persister cells, which fail to induce ATF3 (Fig. 4a), ectopic re-expression of ATF3 decreased STAT1 levels (Fig. 4f). Similarly, rescue of ATF3 expression in DFFB KO persister cells by etoposide treatment also suppressed STAT1 (Fig. 4b). Furthermore, like DFFB LOF cells, ATF3-depleted persister cells are hindered in their ability to regrow into DTEP colonies (Fig. 4g). Also similar to DFFB LOF cells (Extended Data Fig. 4c-p), decreased DTEP colony formation of ATF3-depleted cells was not due to any defect in proliferation or persister cell viability (Extended Data Fig. 8a,b). ATF3 also remained highly expressed in DFFB WT, but not KO, cells at the timepoint WT DTEP colonies form, consistent with a continuous role for ATF3 in suppressing ISGs throughout DTEP colony formation (Extended Data Figs. 7c, 8c).

We next sought to determine how ATF3 suppresses ISG expression in persister cells. ATF3 binds and antagonizes AP1 transcription factor complex proteins (Extended Data Fig. 8d),^49^ and AP1 was recently reported to be drive memory inflammatory gene expression including ISGs in epidermal stem cells.^50^ We therefore tested whether ATF3 suppression of ISGs in persister cells occurs as a result of negative regulation of AP1 transcription factor activity. Consistent with this, we observed that cotreatment of DFFB KO and ATF3-depleted persister cells, which are each deficient in ATF3 and have elevated ISGs (Figs. 3a-c; 4a,d,e and Extended Data Figs. 6l-n and 7a-c), with AP1 inhibitor T-5224 decreased expression of IFN-related gene sets (Fig. 4h,i and Supplementary Tables 12-15). In contrast, AP1 inhibitor treatment has no effect on IFN-related gene sets in WT persister cells as expected for cells with intact ATF3-mediated AP1 inhibition (Extended Data Fig. 8e and Supplementary Tables 16 and 17). These data support ATF3-mediated AP1 inhibition as a mechanism by which DFFB suppresses ISGs in persister cells.

## On-treatment patient tumours exhibit features of IFN-enforced persister cell growth arrest

To assess evidence for our findings in patients, we explored datasets from melanoma and lung cancer patient tumours which were analyzed by RNAseq prior to and during targeted therapy treatment.^51,52^ We first investigated the anastasis signature which marks drug-stressed persister and DTEP cells (Fig. 1b-f and Extended Data Fig. 9a,b) and found that it is increased in both on-treatment melanoma samples and residual disease (RD) lung cancer samples collected during treatment (Extended Data Fig. 9c,d). In contrast, progressive disease (PD) lung cancer samples collected at later timepoints are markedly devoid of the anastasis signature suggesting that PD tumours have acquired resistance and do not experience drug stress (Extended Data Fig. 9d). Indeed, many of the PD tumours were reported to harbor mutations and nongenetic mechanisms which can cause resistance.^49^

Similar to persister and DTEP cells (Extended Data Fig. 6l), both the on-treatment melanoma and RD lung cancer samples have decreased expression of proliferation marker Ki-67 (Extended Data Fig. 9e,f) while PD tumours have high Ki-67 (Extended Data Fig. 9f). On-treatment melanoma and RD lung cancer samples are also enriched for ATF3 and exhibit moderately elevated ISGs (Extended Data Fig. 9g-j) similar to DFFB WT persister and DTEP cells (Fig. 3a and 4a, Extended Data Fig. 6a-c,i,j and 7a-c). In contrast, consistent with a lack of drug stress, PD tumours fail to induce ATF3 (Extended Data Fig. 9j) and exhibit even further elevated ISGs (Extended Data Fig. 9h). Without drug stress to induce tumour cell-intrinsic IFN production, the source of IFN in PD may be primarily from immune infiltrate.^49^ Despite high ISGs, PD tumours are proliferative suggesting that in the absence of drug stress, IFN alone is insufficient to cause tumour growth arrest. Indeed, we also found that elevated STAT1 has minimal effect on proliferation of unstressed parental cells (Extended Data Fig. 6q,r) yet blocks drug stressed persister cell regrowth (Fig. 3j-m). Taken together, these analyses demonstrate that multiple core features of DFFB WT persister and DTEP cells are present within drug stressed residual tumours including anastasis, ATF3, and IFN regulation, but that untreated or fully resistant tumours which avoid drug stress lack these features.

## Discussion

The adaptive mechanisms utilized by cancer cells to survive drug treatment and develop resistance are poorly understood. While genetic resistance mutations often drive long term acquired drug resistance, the earliest events which enable residual cancer cells to regrow may be nongenetic.^53^ By exploring the consequences of sublethal apoptosis in persister cells, we uncovered a regulatory mechanism in which apoptotic DNase DFFB orchestrates transcriptional repression of ISGs, enabling escape from IFN-enforced growth arrest and facilitating initial persister cell regrowth (Figure 4j). Cancer cells deficient in DFFB are strikingly unable to regrow during targeted therapy treatment. DFFB also promotes mutagenesis in persister cells which may contribute to cancer stress-induced mutagenesis.^7,8,29^ These findings are consistent with clinical observations that increased apoptosis signal in regressed tumours during drug treatment predicts worse outcome.^54^

We found DFFB-mediated DNA damage induces ATF3 which accumulates during drug treatment and functions as part of IFN negative feedback machinery that limits persister cell ISG expression. It was recently shown that cancer cells which survive brief treatment with cytotoxic concentrations of BH3 mimetics induce MOMP and cytochrome c release which activates the ISR including ATF4 and its target gene ATF3.^13^ We did not observe ISR activation or ATF4-mediated ATF3 expression in targeted therapy treated persister cells which exhibit lower MOMP and cytochrome c release compared to cells which survive brief cytotoxic BH3 mimetic treatment. Based on these observations we propose that there are distinct outcomes for cancer cells depending on the level and duration of MOMP they experience. Cancer cells which survive transient high levels of MOMP can activate the ISR in a caspase-independent manner to form persister cells and gain metastatic potential,^13^ while targeted therapy treated persister cells experience chronic moderate levels of MOMP and instead undergo caspase-dependent DFFB-mediated ISG suppression which promotes regrowth without affecting persister cell formation. Given these disparate molecular mechanisms and phenotypic consequences of apoptotic stress, it seems likely that there are additional features of sublethal death signaling remaining to be discovered.

While DFFB-deficient persister cells are almost completely unable to regrow due to high levels of IFN signaling, the majority of WT persister cells also remain growth arrested and only a small minority regrow into DTEP colonies. It is unclear why a subset of WT persister cells regrow and most do not. One possibility is heterogeneity in DFFB and IFN signaling levels. Indeed, persister cells with higher levels of caspase activity have higher levels of DFFB-induced DNA damage (Extended Data Fig. 2j) and there is also some heterogeneity in persister cell ISG expression such that DTEP cells that are proliferating have lower ISGs (Fig. 3a-c and Extended Data Figs. 6a-d,i,j,n,o). However, given that ATF3 levels are similar among cycling and noncycling DTEP cells (Extended Data Fig. 8c), additional adaptations likely contribute to DTEP cell cycling which remain to be determined. Beyond IFN suppression, there are other hurdles that persister cells also likely need to overcome to escape quiescence including reestablishment of mitogenic signaling and metabolic processes.^55^ Therefore, while DFFB is a critical factor to enable DTEP formation, a full understanding of the transition from quiescent persister cell into cycling DTEP cell will require additional study.

DFFB is a nonessential gene in mice and a potentially druggable enzyme.^56–58^ Despite being a member of the apoptotic pathway, DFFB is not required for the execution of cell death^59^ as DFFB KO cells respond similarly to drug as WT cells (Extended Data Fig. 4) and display both elevated cleaved caspase 3 (Extended Data Fig. 4q) and anastasis signatures (Extended Data Fig. 9a). In addition to our findings that DFFB-deficient persister cells and residual tumours are unable to regrow during targeted therapy treatment, DFFB-deficient tumours are also sensitized to radiation.^38^ DFFB inhibition may also promote tumour immunity through upregulation of immune-stimulating ISGs such as STING^60^ specifically within drug-stressed tumour cells (Extended Data Fig. 9k,l). Indeed, because DFFB is inactive and bound to DFFA in unstressed cells, it may be possible to achieve DFFB inhibition specifically within tumour cells under targeted therapy or other stress. Therefore, DFFB is a unique adaptation factor leveraged by cancer cells to adapt to drug stress.

## Methods

### Cell lines and culture

A375 (CRL-1619) and BT474 (HTB-20) cells were purchased from the ATCC. PC9 cells were provided by the Altschuler and Wu Lab at UC San Francisco. A375 cells were cultured in DMEM (Thermo Fisher Scientific) supplemented with 10% fetal bovine serum (FBS) and 1% Antimycotic/Antibiotic (AA) (Thermo Fisher Scientific). PC9 cells were cultured in RPMI-1640 medium (Thermo Fisher Scientific) supplemented with 5% FBS and 1% AA. BT474 cells were cultured in RPMI-1640 medium supplemented with 10% FBS and 1% AA. All cell lines were maintained in 5% CO2 atmosphere at 37 °C. Cell lines were split with 0.25% trypsin-EDTA (Thermo Fisher Scientific). Cell line identities were confirmed with STR profiling at the UC Berkeley Cell Culture Facility. All cell lines regularly tested negative for mycoplasma throughout these investigations using the Lonza Mycoalert Mycoplasma Detection Kit.

### Chemical and antibody sources for cell treatments

Dabrafenib (S2807), trametinib (S2673), erlotinib (S7786), osimertinib (S7297), lapatinib (S2111), staurosporine (S1421), puromycin (S9631), doxycycline hyclate (S4163), and Quinoline-Val-Asp-Difluorophenoxymethylketone (QVD) (S7311) were purchased from Selleck Chemicals. Etoposide (E55500) was purchased from Research Products International. JC-1 dye (T3168) and digitonin (AC407565000) were purchased from Thermo Fisher Scientific. Ghost Dyes Violet 510 (13-0870-T100) and Red 780 (13-0865-T100) were purchased from Tonbo Biosciences. Carbonyl cyanide m-chlorophenyl hydrazone (CCCP) was purchased from Abcam (ab141229). BioTracker NucView 530 Red Caspase-3 Dye (PBS) (SCT105) and cytochalasin B (C2743) were purchased from Sigma-Aldrich. JAK inhibitor ruxolitinib (HY-50856), TBK1 inhibitor MRT67307 (HY-13018), thapsigargin (HY-13433), ABT-737 (HY-50907), S63845 (HY-100741), and T-5224 (HY-12270) were purchased from MedChemExpress. Anti-Interferon alpha/beta receptor 1 antibody (ab10739) was purchased from Abcam.

### Drug and chemical treatments

Previously described protocols were utilized.^10,11^ Briefly, A375 cells were treated with 250 nM dabrafenib and 25 nM trametinib for 14 days, PC9 cells were treated with 2.5 μM erlotinib for 10 days, or with 300 nM or 500 nM osimertinib for 9 days, and BT474 cells were treated with 500 nM or 2 μM lapatinib for 10 days to generate persister cells unless otherwise indicated. A375 cells were treated with 250 nM dabrafenib and 25 nM trametinib for 7 weeks for DTEP cell analyses unless otherwise indicated. Pre-derived persister cells were assayed while still in drug unless otherwise noted. In experiments utilizing caspase inhibitor QVD, 10 μM QVD was added together with the respective targeted therapy treatment for the duration of the experiment. Media and drug were replenished every 3-4 days. For etoposide treatment of DFFB KO persister cells, 100 μM etoposide was added to preformed persister cells for 48 hours prior to 24 hours of recovery without any drugs before protein collection. Control DFFB KO persister cells for this experiment (Fig. 4b) similarly received a 24 hour recovery without any drugs prior to protein collection. For thapsigargin treatments, cells were treated with 1 μM thapsigargin for 4 hours. For BH3 mimetic treatments, A375 cells were treated with 5 μM ABT-737 and 10 μM S63845 for 2.5 hours then allowed to recover in media without drugs for 24 hours. PC9 cells were treated with 1.5 μM ABT-737 and 3 μM S63845 for 3 hours then allowed to recover in normal media without drugs for 2 hours.

### DTEP colony formation

Cells were plated at low starting densities to avoid outgrowth of overlapping DTEP colonies. Specifically, A375 cells were seeded at 4,000 cells per 10 cm dish, and 250 nM dabrafenib and 25 nM trametinib media was added the next day. PC9 cells were seeded at 5,000 cells per 10 cm dish, and 2.5 μM erlotinib or 300 nM osimertinib media was added the next day. BT474 cells were seeded at 500,000 cells per 10 cm dish, and 2 μM lapatinib was added 72 hours later. Drug media was refreshed every 3-4 days. At 7 weeks in drug for A375 cells, 5 weeks in drug for PC9 cells, and 11 weeks in drug for BT474 cells, plates were fixed with 90% methanol and stained with crystal violet. The plates were then blinded by a separate laboratory member prior to colony counting by microscopy. Colonies with greater than 25 cells were considered DTEP colonies. To calculate the fraction of cells that regrew into DTEP colonies (“DTEP fraction”), the number of DTEP colonies per plate was divided by the sum of the number of isolated single cells plus colonies of any size.

### Microscopy

Brightfield photos were captured with the EVOS XL Core microscope using the 4X LPlan PH2 objective.

### Cell viability

Cell viability was assessed with CellTiter Glo (CTG) 2.0 Cell Viability assay (Promega, #G7571). Luminescence was read with a Molecular Devices SpectraMax iD3 plate reader with SoftMax Pro 7 software.

### Single cell RNA sequencing

A375 cells were cultured with 250 nM dabrafenib and 25 nM trametinib and PC9 cells were treated with 2.5 μM erlotinib for 2 weeks to establish persister cells. For the DTEP timepoint, A375 WT and DFFB KO cells were cultured with 250 nM dabrafenib and 25 nM trametinib for 9 weeks during which WT cells formed DTEP colonies while KO cells remained alive as persister cells and did not grow into DTEP colonies. For all conditions, an untreated parental control was cultured alongside the drug treated conditions. At the time of analysis, cells were lifted with trypsin and libraries were generated using the 10X Chromium Single Cell 3’ v3 kit (10X Genomics). Quality control of the libraries was conducted with the High Sensitivity D1000 ScreenTape kit with the Agilent TapeStation and then sequenced using a NovaSeq S4 flowcell on an Illumina NovaSeq 6000 sequencer at the UCSD Institute for Genomic Medicine.

### ScRNAseq data processing and mapping

Fastq files were aligned to the human “refdata-cellranger-GRCh38-3.0.0” genome with 10X Genomics Cell Ranger 3.1.0 using the “cellranger count” command to generate single cell feature counts for each library.^61^ The “filtered_feature_bc_matrix” generated for each population was used to create a “Seurat object” in the Seurat R package version 4.0.3.^62–64^ Cells containing greater than 1,000 and less then 7,500 features, and with less than 20% mitochondrial reads were included in downstream analyses. A cell cycle score was calculated for each cell using the default Seurat method with a published list of cell cycle genes,^65^ and this score was used to regress cell cycle during normalization and scaling with the “SCTransform” command. To generate uniform manifold approximation and projection (UMAP) plots, the commands “RunPCA,” “RunUMAP,” “FindNeighbors,” and “FindClusters” were performed with default settings, with 30 dimensions used for “RunUMAP” and “FindNeighbors.” In the “FindClusters” command, the resolution for A375 and PC9 was set to 1.0 and 0.15, respectfully, based on visualization of graphed clusters. For cluster analyses within the persister cell populations only, the resolution was set to 0.2 for A375 and 0.05 for PC9. The Seurat command “FindMarkers” was used without thresholds for fold change or percentage of gene-expressing cells to calculate differentially expressed genes.

### ScRNAseq persister cell gene signature analysis

Gene set enrichment analysis was conducted with the ClusterProfiler R package (version 3.18.0) to calculate a normalized enrichment score (NES) for each gene set in the specified population.^66^ The minimum gene set size for analysis was set to five and the Benjamini-Hochberg method was used for false discovery rate correction. Signature scoring was performed using the Seurat command “AddModuleScore.” To display the enriched Hallmarks interferon alpha response gene set on a UMAP, AUCell package version 1.12.0 was to calculate gene set scores per cell.^67^

### JC-1 mitochondrial membrane potential assay

A375 persister cells were derived with 250 nM dabrafenib and 25 nM trametinib for 2 weeks and untreated parental cells were cultured alongside. Cells were then lifted with trypsin for JC-1 staining. As a positive control, cells were treated with 50 µM carbonyl cyanide m-chlorophenyl hydrazone (CCCP) for 5 minutes. JC-1 was dissolved in DMSO (Thermo Fisher Scientific, #D12345) at a final concentration of 1.5 µM and cells were stained for 30 minutes away from light at 37 °C. Cells were then stained with Ghost Dye Red 780 (Tonbo Biosciences, #13-0865-T100) diluted 1:1000 in PBS (Gibco, #10010023) for 15 minutes away from light at room temperature. Cells were washed in PBS and analyzed on a BD FACSCanto RUO flow cytometer using 488 nm and 640 nm lasers for green and red fluorescence respectively. At least 30,000 live cell events were collected per sample. The flow cytometry results were analyzed using FlowJo Software (BD Life Sciences) version 10.7.1. See Supplementary Fig. 3 for gating strategy.

### Immunoblotting

Persister or DTEP cells were derived with drug treatment in 10 or 15 cm plates. Cells were then washed with PBS and lysed using RIPA buffer (Thermo Fisher Scientific, #89900) supplemented with phosphatase inhibitor (Thermo Fisher Scientific, #78420) and protease inhibitor (Thermo Fisher Scientific, #78430). Lysates were centrifuged at 14,000 g at 4 °C for 15 min, and the protein concentration of the supernatant was determined using the Pierce BCA Protein Assay Kit (Thermo Fisher Scientific, #23225). Lysates were mixed with sample buffer (Thermo Fisher Scientific, NP0007) and denatured at 70 °C for 10 min. Samples were separated by SDS–PAGE (Bolt 4–12% Bis-Tris Gel, Life Technologies, NW04120BOX), run with Chameleon Duo Pre-stained Protein Ladder (LICOR, #928-60000), and transferred to a nitrocellulose membrane using the iBLOT 2 Dry Blotting System (Life Technologies, IB21001). Membranes were blocked with 5% BSA for 1 hour at room temperature, and then incubated with primary antibody at 4 °C overnight. LICOR secondary antibodies were then incubated with the membrane for 1 hour at room temperature, and membranes were imaged using the LICOR Odyssey Imaging System and Image Studio version 5.2. β-Tubulin or vinculin levels were measured as a loading control. Antibody commercial sources were: β-Tubulin (Invitrogen, MA5-16308); Vinculin (Cell Signaling Technology, #4650); γH2AX (Cell Signaling Technology, #9718); Cleaved caspase 3 (Cell Signaling Technology, #9664); STAT1 (Cell Signaling Technology, #9176); phosphorylated STAT1 (Cell Signaling Technology, #9167); phosphorylated TBK1 (Cell Signaling Technology, #5483); phosphorylated IRF3 (Cell Signaling Technology, #37829); ATF3 (Cell Signaling Technology, #33593); STING (Cell Signaling Technology, #13647); Caspase 9 (Cell Signaling Technology, #9502); Cytochrome c (BioLegend, #612310); DFFB (Santa Cruz Biotechnology, SC-374067); DFFA (Abcam, ab108521); ATF4 (Cell Signaling Technology, #11815); phosphorylated eIF2α (Cell Signaling Technology, #3398); eIF2α (Cell Signaling Technology, #5324); IRDye 680RD Goat anti-Mouse IgG secondary antibody (LICOR, #926-68070); IRDye 800CW Goat anti-Mouse IgG secondary antibody (LICOR, #926-32210); IRDye 680RD Goat anti-Rabbit IgG secondary antibody (LICOR, #926-68071); IRDye 800CW Goat anti-Rabbit IgG secondary antibody (LICOR, #926-32211).

### DNA damage, cleaved caspase 3, and cytochrome c flow cytometry

A375 persister cells were derived with 250 nM dabrafenib and 25 nM trametinib for 2 weeks and untreated parental cells were cultured alongside. For γH2AX experiments, A375 persister cells treated with 10 µM QVD for the duration of drug treatment were also prepared. After 2 weeks of culture, cells were trypsinized and collected for staining. Cells were stained with viability dye Ghost Dye Red 510 or 780 diluted 1:1000 in PBS for 15 minutes away from light at room temperature. For cytochrome c staining, incubation in 50 µg/mL digitonin in PBS for 10 minutes on ice was performed to selectively permeabilize the cell membrane.^68^ Cells were fixed with 4% paraformaldehyde for 10 minutes at room temperature. For γH2AX and cleaved caspase 3 staining, cells were permeabilized with 0.3% Triton X-100 in PBS for 10 minutes at room temperature. Cells were stained for proteins of interest using primary conjugated antibodies at manufacturer recommended dilutions in PBS for 30 minutes to 1 hour at room temperature. Cells were washed in PBS and analyzed on a BD FACSCanto RUO flow cytometer using 405 nm and 640 nm lasers for blue and red fluorescence respectively. At least 30,000 live cell events were collected per sample. Flow cytometry data were analyzed using FlowJo Software (BD Life Sciences) version 10.7.1. See Supplementary Figs. 2 and 7 for cytochrome c flow cytometry, Supplementary Fig. 4 for cleaved caspase 3 gating strategy, and Supplementary Fig. 6 for γH2AX gating strategy.

### Caspase 3/7 activity reporter assay

Caspase activity was measured using NucView 530 Caspase-3 Substrate (Biotium #10408). A375 persister cells were pre-derived from 2 week treatment with 250 nM dabrafenib with 25 nM trametinib while parental cells were cultured alongside. Positive control cells were derived from 4-hour parental cell treatment with 1 µM staurosporine and these pre-apoptotic cells were assayed by flow cytometry prior to death. Adherent tumour cells were collected by trypsinization (∼1 million cells/sample), washed in PBS, and incubated with Ghost Dye Violet 510 cell viability dye following the manufacturer’s instructions. Cells were washed with PBS + 1% FBS and subsequently incubated with 5 µM NucView 530 Caspase-3 Substrate for 30 minutes in PBS + 1% FBS and analyzed by flow cytometry using a BD FACSAria II sorter. Cells not treated with NucView 530 Caspase-3 Substrate were used as unstained controls. Fluorescence was measured as follows: Ghost Dye Violet 510 (405 nm laser, 525/50 filter), NucView 530 Caspase-3 Substrate (488 laser, 530/30 filter). See Supplementary Fig. 5 for gating strategy. Sorted live cells were collected in PBS + 10% FBS and used for downstream viability assays. For testing caspase 3/7 activity-positive persister cell regrowth, cells were sorted and plated at 1,000 cells per well in 12-well plates in drug-free media. 24 hours later, CTG was performed to measure cell viability. To assess regrowth ability, drug-free media was refreshed on day 3 and CTG was performed on day 6. The DNA damage levels for A375 persister cells with basal and medium caspase activity were measured by sorting cells and extracting protein for western blot analysis as described above.

### CRISPR editing

Caspase 9 knockout cells were generated with Santa Cruz caspase 9 CRISPR plasmids (h) (cat#: sc-400257-KO-2), ATF3 knockout cells with Santa Cruz ATF3 CRISPR plasmids (h) (cat#: sc-416577), and ATF4 knockout cells with Santa Cruz ATF4 CRISPR plasmids (h2) (cat#: sc-400155-KO-2) following manufacturer’s protocols. Briefly, caspase 9 CRISPR plasmid was co-transfected with caspase 9 HDR plasmid (h2) (cat#: sc-400257-HDR-2), ATF3 CRISPR plasmid was co-transfected with ATF3 HDR plasmid (h) (cat#: sc-416577-HDR), and ATF4 CRISPR plasmid was co-transfected with ATF4 HDR plasmid (h2) (cat#: sc-400155-HDR-2) using UltraCruz Transfection Reagent (cat#: sc-395739) into A375 or PC9 cells. Cells were then selected with puromycin, expanded, and edited cell pools were analyzed for depletion of target protein via western blot. Caspase 9 (Extended Data Figs. 2i and 3g), ATF3 (Fig. 4g), and ATF4 (Extended Data Fig. 7l,m) were strongly depleted. ATF3 depletion was assessed in persister cells because ATF3 is not expressed in parental cells while ATF4 was assessed in cells treated with thapsigargin to induce ATF4 expression. Caspase 9-depleted cells also underwent clonal selection to identify cells with complete KO, and the depleted cells (pooled KO) or the clonal complete KO clones were utilized in experiments as described. The ATF3- and ATF4-depleted (pooled KO) cell pools were used without further clonal selection. DFFB CRISPR editing was performed at the UC San Francisco Cell and Genome Engineering Core. Human DFFB sgRNA g1 was previously reported.^18^ Additional guide RNAs targeting the first (g4) or second (g2r) exon of human DFFB were designed and cloned into the eSpCas9(1.1) vector from the Zhang laboratory at MIT (Addgene plasmid #71814; RRID:Addgene_71814). A375 DFFB KO clone #1 was constructed with sgRNA g1, A375 DFFB KO clone #2 was constructed with g4, PC9 DFFB KO clones were both constructed with g2r, and BT474 DFFB KO clones were both constructed with g4.

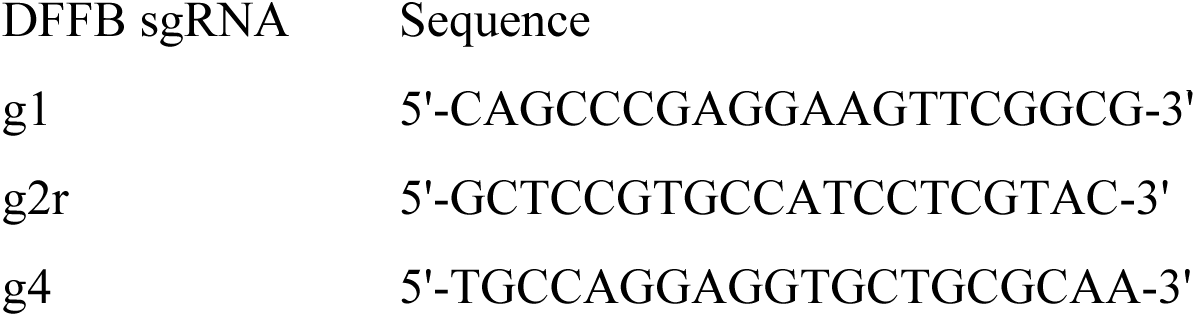

Following sequence verification, the guide constructs were introduced individually either by transfection or electroporation. For transfection, guide constructs were co-transfected with a vector expressing mCherry and puromycin resistance using Lipofectamine 3000. Transfected cells were selected with 1 µg/mL puromycin for 3-4 days. For electroporation, cells were electroporated with the CRISPR/Cas9 RNP gRNAs and high-fidelity Cas9p (Integrated DNA Technologies) using the Lonza 4D Nucleofector kit SF with program FF-120. After cell expansion, genomic DNA was extracted from a portion of the cell pool (Nucleospin Blood gDNA extraction kit, Takara Bio) and the target sites were amplified (Phusion DNA polymerase, NEB) and sequenced using the primers indicated in the table below (Sanger sequencing by GeneWiz, South San Francisco, CA; primers and sgRNA cloning oligonucleotides from Integrated DNA Technologies). CRISPR/Cas9 efficiency was determined by TIDE analysis (tide.deskgen.com). Clones which showed only frameshift-causing indels in all alleles were further analyzed by TOPO cloning (Life Technologies) and Sanger sequencing and with the ICE analysis tool (Synthego). Clones with a DFFB KO genotype were subsequently confirmed to lack DFFB protein expression by western blot (Extended Data Fig. 3d-f).

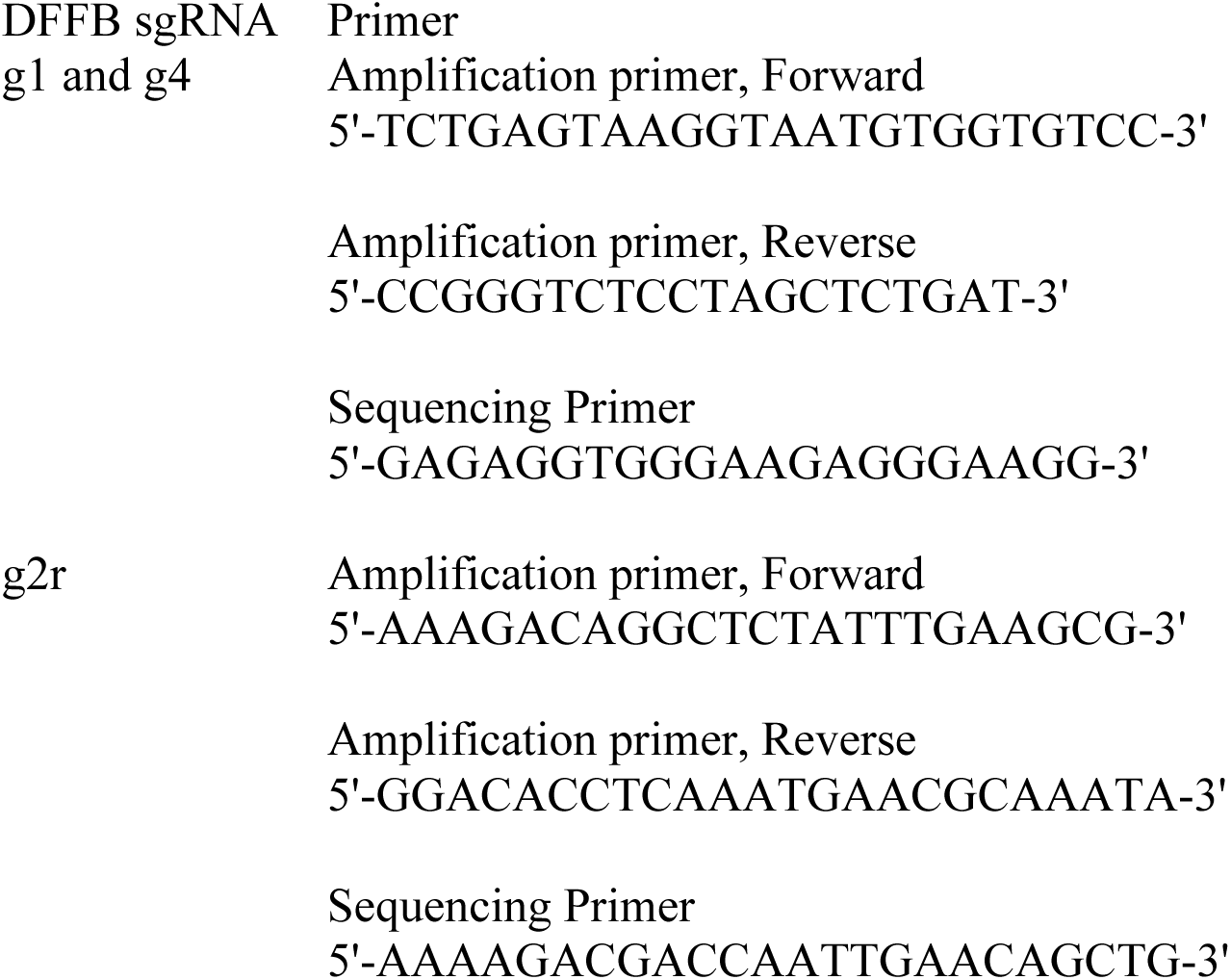

### Lentivirus production

E. coli cultures with lentiviral vectors were grown on LB Agar Ampicillin-100 plates (Sigma-Aldrich, #L5667). A colony was picked and grown overnight in LB-amp and plasmid DNA was isolated using the Qiagen HiSpeed Plasmid Midi Kit (Qiagen #12643) and quantified with a Nanodrop. HEK293T cells were transfected with packaging plasmids psPAX2 (Addgene, #12260) and pMD2G (Addgene, #12259) in Opti-MEM (Thermo Fisher Scientific, #31985062) and polyethylenimine (Sigma-Aldrich, #764604). Virus-containing supernatant was collected, centrifuged, then filtered with a 0.45 μm filter (Sigma-Aldrich, #SE1M003M00).

### Ectopic gene expression

Doxycycline-inducible DFFB expression vectors, STAT1 overexpression vector, and ATF3 vector were obtained from VectorBuilder. The cleavage resistant DFFA (DFFA-CR) short isoform construct was provided by the Elledge lab.^25^ DFFB expression was driven by the 3^rd^ generation tetracycline-responsive element promoter (TRE3G), STAT1 expression was driven by the ubiquitin C promoter (UBC), and ATF3 expression was driven by the cytomegalovirus immediate early enhancer/promoter (CMV). All vectors contained a puromycin resistance gene driven by the phosphoglycerate kinase 1 promoter. Lentivirus was produced as described above. The DFFA-CR construct was transduced into A375 WT cells, STAT1 overexpression construct into A375 WT and PC9 WT cells, and WT DFFB [NM_004402.4], nuclease-deficient DFFB (H260N),^26^ or human ATF3 into A375 DFFB KO cells. Cells were selected with 2 μg/mL puromycin for 3 days. A single cell clone confirmed for DFFA-CR expression was isolated and expanded for experiments. For doxycycline-inducible DFFB, cells were seeded and 1.0 μg/mL doxycycline was added the following day. Media with doxycycline was refreshed every 3-4 days for the duration of the experiment.

### Whole exome sequencing

To eliminate preexisting genetic heterogeneity, a single A375 or PC9 WT or DFFB KO cell was used to start the experiments. Following initial expansion for 3 weeks, a portion of the cell population was frozen for the reference sequence and the remaining cells were plated onto 10 cm dishes at 4,000 cells per dish. 250 nM dabrafenib plus 25 nM trametinib or 2.5 μM erlotinib was added the following day, and drug media was refreshed every 3-4 days. Following 7 weeks of drug treatment for A375 and 10 weeks treatment for PC9, the DFFB WT plates were composed of primarily DTEP colonies while the DFFB KO plates were almost exclusively isolated cells or small clumps of surviving persister cells which had not regrown (see Fig. 2g,h). Then, each plate was allowed to regrow without drug for 10 days, cells were lifted with trypsin, and multiple individual cells were isolated through single cell bottlenecks in 384 well plate wells to purify their genotypes. The isolated single cells were subsequently expanded for 5-6 weeks in drug-free media in increasing well sizes up to a 15 cm plate to accumulate adequate genomic DNA content for genomic sequencing. Frozen cell pellets had DNA extracted and libraries prepared for whole exome sequencing (Agilent SureSelect V6 58M) or whole genome sequencing (A375 pre-treatment reference sample, NEB Ultra II) at Novogene and were sequenced on an Illumina NovaSeq 6000 PE150.

### Single nucleotide variant analysis

Each sequenced sample was analyzed according to the GATK best practices workflow for data preprocessing for variant discovery followed by the somatic short variant discovery workflow.^69–71^ GATK version 4.2.4.0 pipelines were used. Fastq files were mapped to the hg38 reference genome provided in the GATK shared resource bucket with BWA.^72^ Duplicate reads were marked with Picard, and base scores were recalibrated with the GATK tool “BaseRecalibrator.” Mutect2 was used to call mutations between the drug treated sample and the respective starting population reference sample. Sequencing errors and contamination were estimated with “GetPileUpSummaries”, “CalculateContamination”, and “FilterMutectCalls”. Variants were annotated with “Funcotator”. Low frequency (< 0.4) mutations, which could arise post-treatment during expansion from isolated single cells prior to sequencing, were excluded from the analysis. Output files were analyzed with the Maftools R Bioconductor package version 2.6.05.^73^

### In vivo tumour acquired resistance

The University of California San Francisco Institutional Animal Care and Use Committee (IACUC) approved the xenograft studies which were performed at the UCSF Preclinical Therapeutics Core in protocol #AN179937. Sample size was determined based on prior experience with this model.^11^ A375 cells were cultured in DMEM with 10% FBS prior to implantation. Cells were suspended in a 1:1 mixture of PBS:matrigel to a final concentration of 1 x 10^8^ cells/mL. 10 million DFFB WT and KO A375 cells were subcutaneously injected into opposing flanks of 6-8 week old female NSG mice (Jackson Laboratory, #005557). When tumour volumes reached 100-200 mm^3^, drug dosing was started via once daily oral gavage with 100 mg/kg dabrafenib and 1 mg/kg trametinib. Tumour volumes were measured 1-2 times per week by blinded observers. Tumour volumes were normalized to the minimum volume of each tumour during treatment and fold change of tumour regrowth from this minimum volume was calculated. To focus analysis on acquired resistance, only mice with at least one relapsed tumour (WT or KO), defined by 10-fold regrowth from the minimal volume during treatment, were included in Fig. 2l. Additional mice which did not exhibit tumour relapse of either WT or KO tumours were excluded from Fig. 2l but are included in the Fig. 2 source data.

### RNA extraction and quantitative RT-PCR

RNA from A375 parental cells and cells treated with 1 μM dabrafenib and 100 nM trametinib with and without 10 μM QVD for 6 days was extracted with TRIzol (Thermo Fisher #15596026) following manufacturer’s protocol. RNA was converted to cDNA with the RevertAid First Strand cDNA Synthesis Kit (Thermo Fisher #K1621) prior to quantifying IFNB1 levels with TaqMan assay (Thermo Fisher #4331182).

### Bulk RNA sequencing analysis

Biological triplicates of A375 WT, DFFB KO cl1, and ATF3-depleted (pooled KO) parental cells, persister cells derived from 14-day treatment with 250 nM dabrafenib and 25 nM trametinib, and persister cells co-treated for the duration of persister formation with 20 μM AP1 inhibitor T-5224 were trypsinized and RNA was isolated with the RNeasy Mini Kit (Qiagen, 74104). The libraries were constructed with Illumina Stranded mRNA Prep and sequenced using a NovaSeq S4 flowcell on an Illumina NovaSeq 6000 sequencer at the UCSD Institute for Genomic Medicine. Analysis was performed using *Partek Flow* software (v12.1.0) with default options unless otherwise noted. Briefly, bases were trimmed based on quality score (Phred > 35) from the 3’ end. Reads were aligned to human hg38 reference genome with STAR aligner, v2.7.8a. Then, reads were quantified with the *Partek* task quantify to annotation model using the Ensembl 112 release. Differentially expressed genes were calculated with DESeq2 using default settings (v1.46.0) and gene set enrichment analysis was performed with clusterProfiler (v4.14.6) after removing genes with an adjusted p-value equal to “NA”. The Benjamini-Hochberg method was used for false discovery rate correction unless otherwise indicated.

### Cytosolic mitochondrial DNA isolation and PCR

Cytosolic DNA was isolated as previously described.^74^ Briefly, cells from each condition were split into two pellets. For the whole cell fraction, DNA was extracted from the first pellet with the DNeasy Blood and Tissue Kit (Qiagen #69504). For the cytosolic fraction, the second pellet was resuspended in 150 mM NaCl, 50 mM HEPES pH 7.4, and 25 μg/mL digitonin for 10 minutes at room temperature. This mixture was centrifuged (150 x *g*) at 4 °C for 10 minutes and supernatant serially collected three times. The supernatant was then centrifuged (17,000 x *g*) at 4 °C for 10 minutes and the pellet DNA was cleaned with the Qiaquick Nucleotide Removal Kit (Qiagen #28115). All DNA was quantified by spectrophotometry. The following primers were used for PCR with PowerTrack SYBR Green Master Mix for qPCR (ThermoFisher, A46012): hMT-Dloop

Forward: CATCTGGTTCCTACTTCAGGG

Reverse: CCGTGAGTGGTTAATAGGGTG

Cytosolic fraction measurements were normalized to the respective whole cell fraction measurement.

## Data availability

Single cell and bulk RNA sequencing data have been deposited in NCBI’s Gene Expression Omnibus (accession number: GSE196018). Whole exome sequencing and whole genome sequencing data have been deposited in the NCBI Sequence Read Archive (PRJNA800470).

## Statistical analyses

All statistical tests were performed with GraphPad Prism version 9.0.0 (86), R version 4.0.3, or RStudio version 1.2.5033. For P values, ns indicates P > 0.05. Differentially expressed (DE) genes from scRNAseq were calculated using the Wilcoxon Rank Sum test. DE genes from bulk RNA sequencing data were calculated using the Wald test in DESeq2. Signature scoring significance between selected populations was calculated with the Mann-Whitney test. Multiple testing correction was performed using the Benjamini-Hochberg method.

## Acknowledgements

We acknowledge Jean Y. J. Wang (UCSD), Richard Gallo (UCSD), Dong-er Zhang (UCSD), Steven Altschuler (UCSF), and Lani Wu (UCSF) for useful discussions. This publication includes data generated at the UC San Diego Institute for Genomic Medicine Genomics Center utilizing an Illumina NovaSeq 6000 that was purchased with funding from a National Institutes of Health SIG grant (#S10 OD026929). This work was performed with the support of the Flow Cytometry Core at the San Diego Center for AIDS Research (P30 AI036214), the VA San Diego Health Care System, and the San Diego Veterans Medical Research Foundation. This research was supported by fellowship support for A.F.W. from the UCSD Cancer Biology, Informatics & Omics T32 training grant CA067754 and the Contemporary Approaches to Cancer Cell Signaling and Communication T32 training grant CA009523 and by UCSD Moores Cancer Center American Cancer Society Institutional Research Grant IRG-15-172-45, NIH/NCI R01CA212767, V Foundation for Cancer Research V Scholar Award V2021-035, Bristol-Myers Squibb – Melanoma Research Alliance Young Investigator Award 689282, University of California Cancer Research Coordinating Committee Award C23CR5537, American Cancer Society IRG Grant # IRG-19-230-48-IRG, UC San Diego Moores Cancer Center Specialized Cancer Center Support Grant NIH/NCI P30CA023100, University of California Academic Senate Bridge Grant BG104446, CDMRP MRP Idea Award ME220037, and CDMRP MRP MASA ME230211 (M.J.H.).

## Author contributions

A.F.W. and M.J.H. conceived and supervised the study and wrote the manuscript; A.F.W., D.A.G.G., C.E.T., M.H., M.X.W., A.E.S., A.H.W., B.E.M., M.H.P., and C.P.L. performed the experiments and analyzed the data; J.L.P. and S.H.H. generated CRISPR KO cell lines; A.F.W. performed bioinformatic analyses.

## Data availability

All data are available from the authors without restriction. Single cell and bulk RNA sequencing data have been deposited in NCBI’s Gene Expression Omnibus (accession number: GSE196018). Whole exome and whole genome sequencing data have been deposited in the NCBI Sequence Read Archive (PRJNA800470).

## Supplementary information

Supplementary Figure 1, uncropped western blot images

Supplementary Figure 2, mitochondrial release of cytochrome c flow cytometry assay schematic

Supplementary Figure 3, loss of mitochondrial potential flow cytometry assay schematic

Supplementary Figure 4, cleaved caspase 3 flow cytometry schematic

Supplementary Figure 5, caspase 3/7 activity reporter flow cytometry schematic

Supplementary Figure 6, γH2AX DNA damage flow cytometry schematic

Supplementary Figure 7, parental, BH3-treated cell, and persister cell mitochondrial release of cytochrome c flow cytometry assay schematic

Supplementary Table 1, differentially expressed genes from bulk RNAseq of A375 WT persister versus parental cells

Supplementary Table 2, enriched Hallmarks gene sets from bulk RNAseq of A375 WT persister versus parental cells

Supplementary Table 3, genes in the anastasis gene set

Supplementary Table 4, whole exome sequencing acquired mutation list from drug treated A375 WT and DFFB KO cells

Supplementary Table 5, whole exome sequencing acquired mutation list from drug treated PC9 WT and DFFB KO cells

Supplementary Table 6, differentially expressed genes from scRNAseq of A375 DFFB WT versus KO cells at the persister and DTEP timepoints

Supplementary Table 7, enriched Hallmarks gene sets from scRNAseq of A375 DFFB WT versus KO cells at the persister and DTEP timepoints

Supplementary Table 8, differentially expressed genes from bulk RNAseq of A375 DFFB KO versus WT persister cells

Supplementary Table 9, enriched Hallmarks gene sets from bulk RNAseq of A375 DFFB KO versus WT persister cells

Supplementary Table 10, differentially expressed genes from bulk RNAseq of A375 ATF3-depleted versus WT persister cells

Supplementary Table 11, enriched Hallmarks gene sets from bulk RNAseq of A375 ATF3-depleted versus WT persister cells

Supplementary Table 12, differentially expressed genes from bulk RNAseq of A375 ATF3-depleted persister cells treated with versus without AP1 inhibitor

Supplementary Table 13, enriched Hallmarks gene sets from bulk RNAseq of A375 ATF3-depleted persister cells treated with versus without AP1 inhibitor

Supplementary Table 14, differentially expressed genes from bulk RNAseq of A375 DFFB KO persister cells treated with versus without AP1 inhibitor

Supplementary Table 15, enriched Hallmarks gene sets from bulk RNAseq of A375 DFFB KO persister cells treated with versus without AP1 inhibitor

Supplementary Table 16, differentially expressed genes from bulk RNAseq of A375 WT persister cells treated with versus without AP1 inhibitor

Supplementary Table 17, enriched Hallmarks gene sets from bulk RNAseq of A375 WT persister cells treated with versus without AP1 inhibitor

## Materials & correspondence

Correspondence and request for materials should be addressed to Matthew J. Hangauer.

---

**Extended Data Fig. 1:**
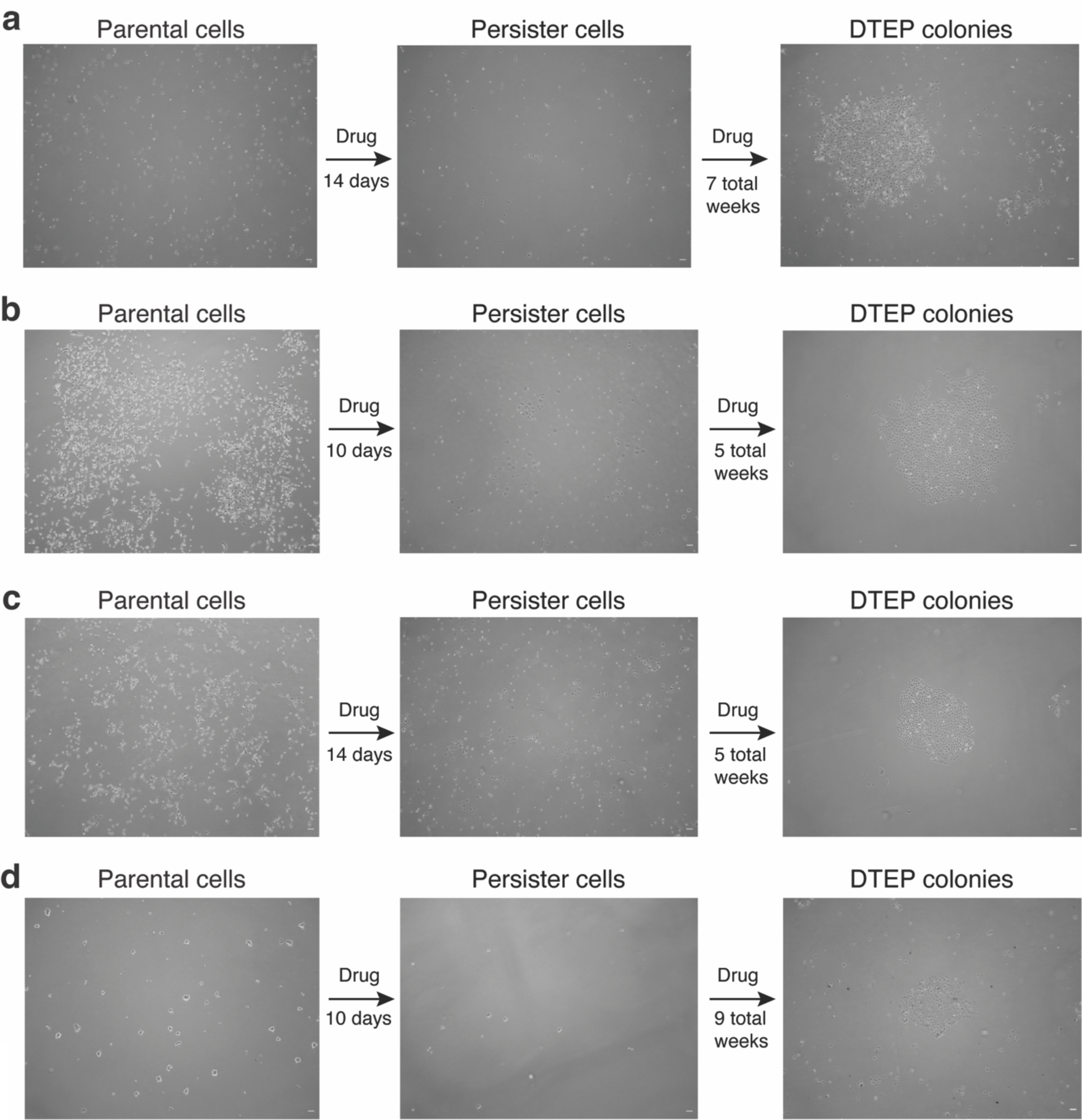
Persister cell and DTEP colony formation models. **a**, A375 BRAF V600E melanoma cells treated with 250 nM dabrafenib and 25 nM trametinib. **b**, PC9 EGFR mutant lung adenocarcinoma cells treated with 2.5 μM erlotinib. **c**, PC9 cells treated with 500 nM osimertinib. **d**, BT474 HER2+ breast cancer cells treated with 2 μM lapatinib. Time shown to DTEP colony formation includes initial persister cell formation treatment period. Scale bar = 100 μm.

**Extended Data Fig. 2:**
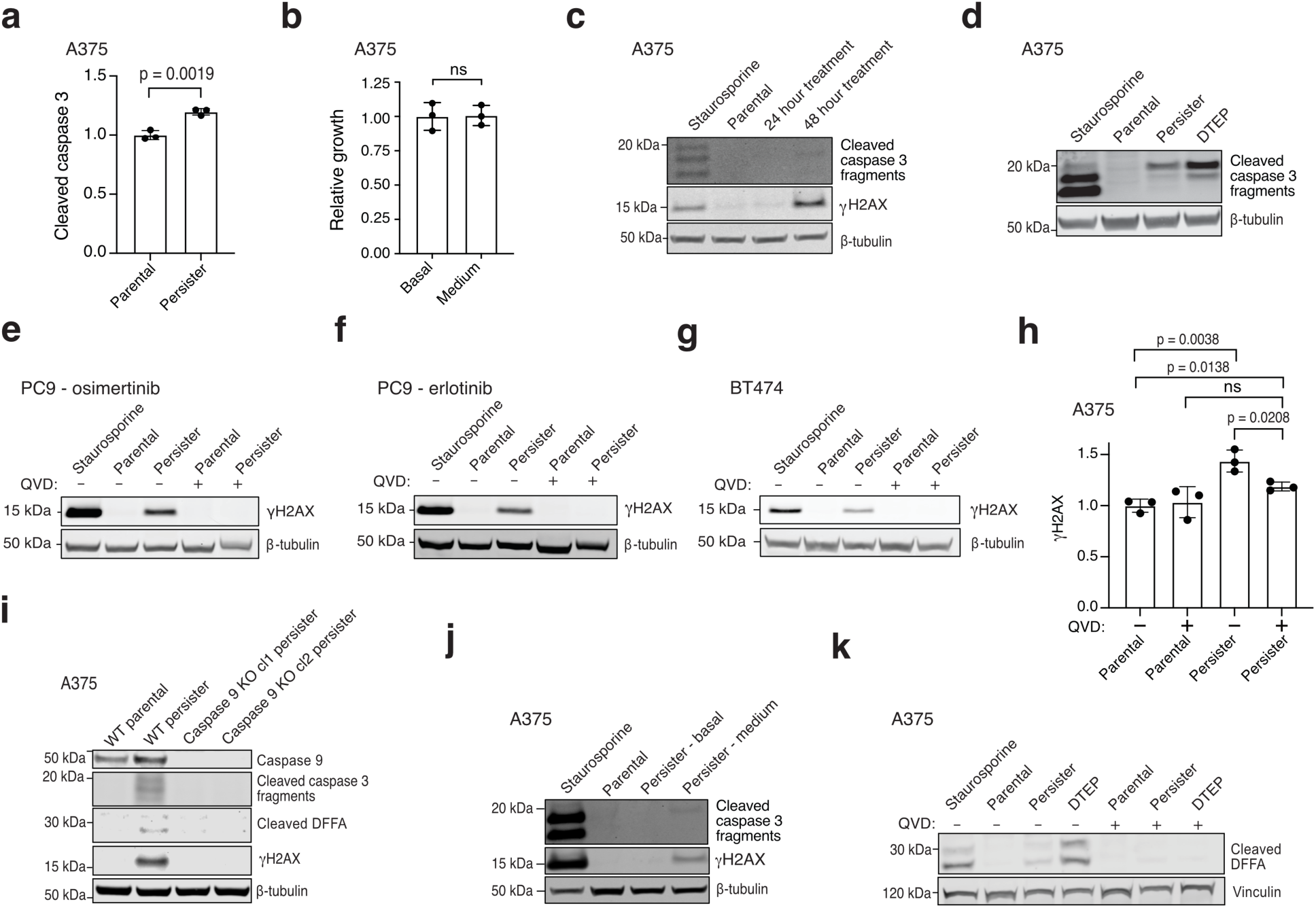
Persister cells undergo chronic drug-induced sublethal apoptotic stress that causes DNA damage. **a**, Quantification of cleaved caspase 3 from flow cytometry of live-gated cells. **b**, Relative growth in drug-free media for 6 days of A375 persister cells after sorting for basal or medium (sublethal elevated) caspase 3/7 activity. See Supplementary Fig. 5 for sorting strategy. **c**, A375 cells treated with 250 nM dabrafenib and 25 nM trametinib for 24 or 48 hours and assessed for cleaved caspase 3 or DNA damage marker γH2AX. **d**, A375 parental, persister, and DTEP cells derived from 7 weeks of treatment with 250 nM dabrafenib and 25 nM trametinib analyzed for cleaved caspase 3. **e-h**, Parental and persister cells assessed for γH2AX with and without treatment with 10 μM caspase inhibitor QVD. **e**, PC9 cells treated with 500 nM osimertinib. **f**, PC9 cells treated with 2.5 μM erlotinib. **g**, BT474 cells treated with 500 nM lapatinib. **h**, A375 parental and persister cells assessed by flow cytometry for γH2AX. **i**, A375 WT and caspase 9 KO clones analyzed for caspase 9, cleaved caspase 3, cleaved DFFA, and γH2AX. **j**, A375 persister cells sorted by caspase 3/7 activity levels and assessed for cleaved caspase 3 and γH2AX. **k**, A375 persister and DTEP cell caspase-dependent cleaved DFFA. 250 nM (**c**,**d**,**j**,**k**), 500 nM (**e**,**f**), or 750 nM (**g**) staurosporine treatment for 24 hours was used as a positive control. For **a**,**b**,**h**, *n* = 3 biological replicates, mean ± s.d.; two-tailed Student’s t-test; ns, not significant.

**Extended Data Fig. 3:**
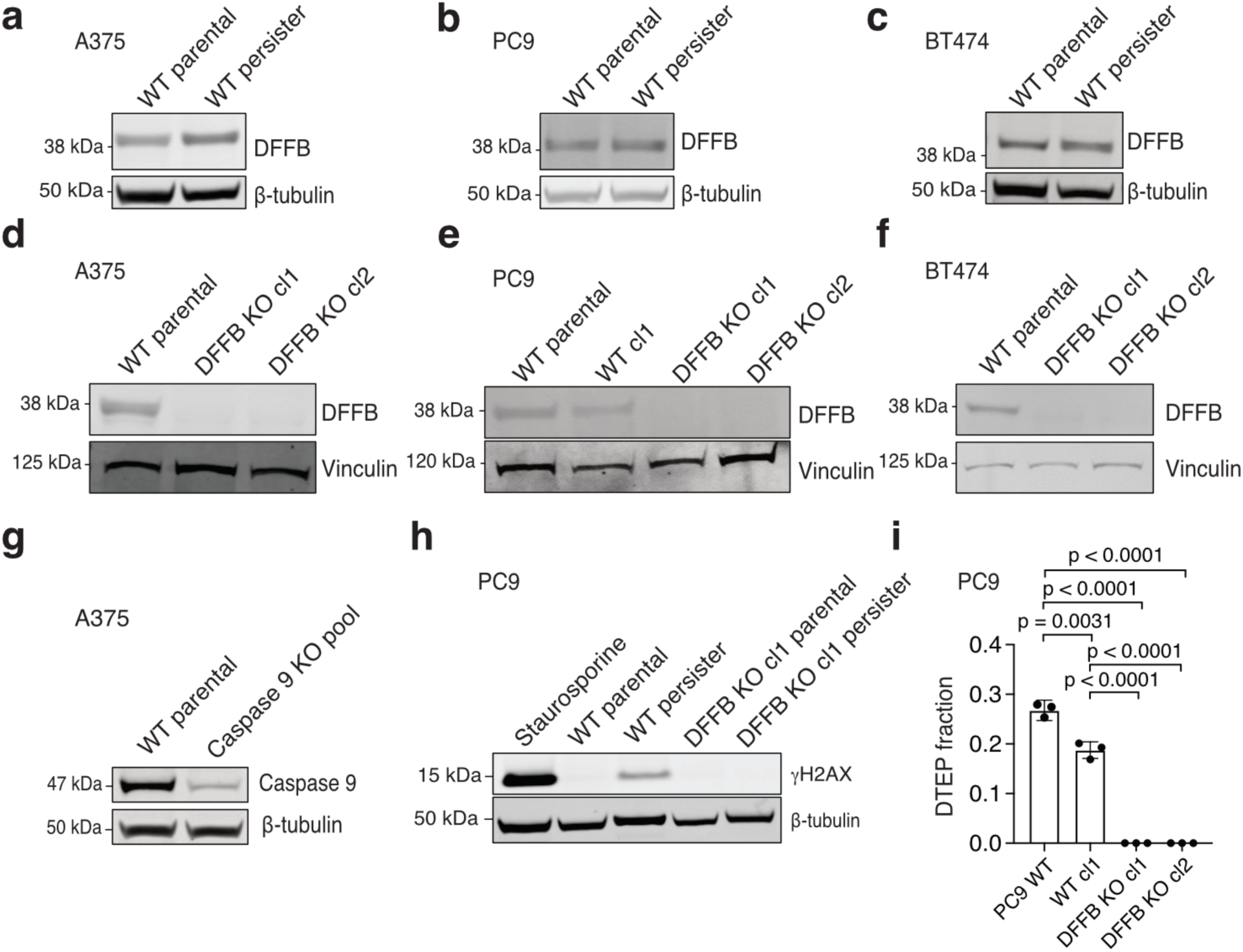
CRISPR KO, DNA damage and DTEP formation data. **a-c**, DFFB expression in parental and persister cells. **a**, A375 cells treated with 250 nM dabrafenib and 25 nM trametinib. **b**, PC9 cells treated with 2.5 μM erlotinib. **c**, BT474 cells treated with 500 nM lapatinib. **d-f**, DFFB loss of expression in CRISPR KO clones. cl, clone. **d**, A375 DFFB KO clones. **e**, PC9 DFFB KO clones. WT cl1 is a control WT clone which failed to lose DFFB expression during CRISPR editing. **f**, BT474 DFFB KO clones. **g**, Caspase 9 expression in CRISPR-mediated caspase 9-depleted (pooled KO) A375 cells used in Fig. 3f. **h**, PC9 cells treated with 2.5 μM erlotinib were assessed for DNA damage marker γH2AX. Parental cells were treated for 24 hours with 500 nM staurosporine as a positive control. **i**, PC9 cells treated with 2.5 μM erlotinib for 5 weeks were analyzed for DTEP formation. *n* = 3 biological replicates, mean ± s.d.; two-tailed Student’s t-test.

**Extended Data Fig. 4:**
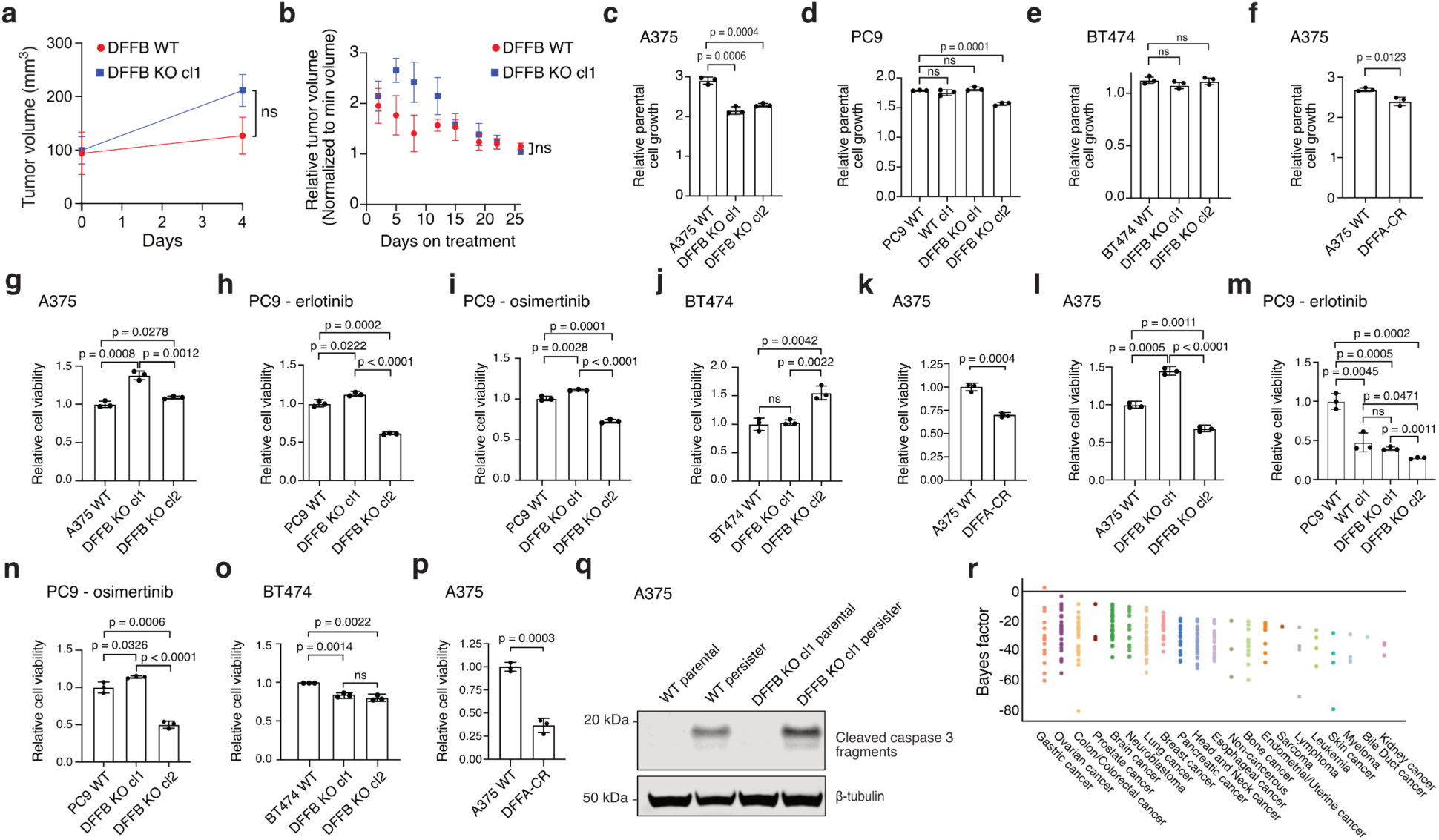
DFFB deficiency does not correlate with clonal variability in tumour cell proliferation or initial drug response. **a-b**, A375 DFFB KO cells form tumours and respond to drug in vivo similarly to WT cells. cl, clone. **a**, A375 xenograft tumour formation and **b,** initial drug response to dabrafenib and trametinib in mice. n = 6 biological replicates; mean ± s.d. is shown; P values calculated with two-tailed Student’s t-test; ns, not significant. **c-f**, Clonal variability in parental cell growth rates in cell culture is independent of DFFB WT or loss of function (LOF) status. DFFA-CR, cleavage resistant DFFA. **g-k**, WT and DFFB LOF cell initial 3 day drug treatment response. **g**,**k**, A375 cells treated with 250 nM dabrafenib and 25 nM trametinib. **h**, PC9 cells treated with 2.5 μM erlotinib. **i**, PC9 cells treated with 500 nM osimertinib. **j**, BT474 cells treated with 2 μM lapatinib. **l-p**, WT and DFFB LOF persister cell formation. **l**,**p**, A375 cells treated with dabrafenib and trametinib. **m**, PC9 cells treated with erlotinib. **n**, PC9 cells treated with osimertinib. **o,** BT474 cells treated with lapatinib. For **c-p**, *n* = 3 biological replicates; mean ± s.d. is shown; P values calculated with two-tailed Student’s t-test. ns, not significant. **q**, Cleaved caspase 3 analyzed in A375 WT and DFFB KO parental and persister cells. **r**, DFFB essentiality in CRISPR screen data from the PICKLES database.^75^ Negative scores reflect non-essentiality.

**Extended Data Fig. 5:**
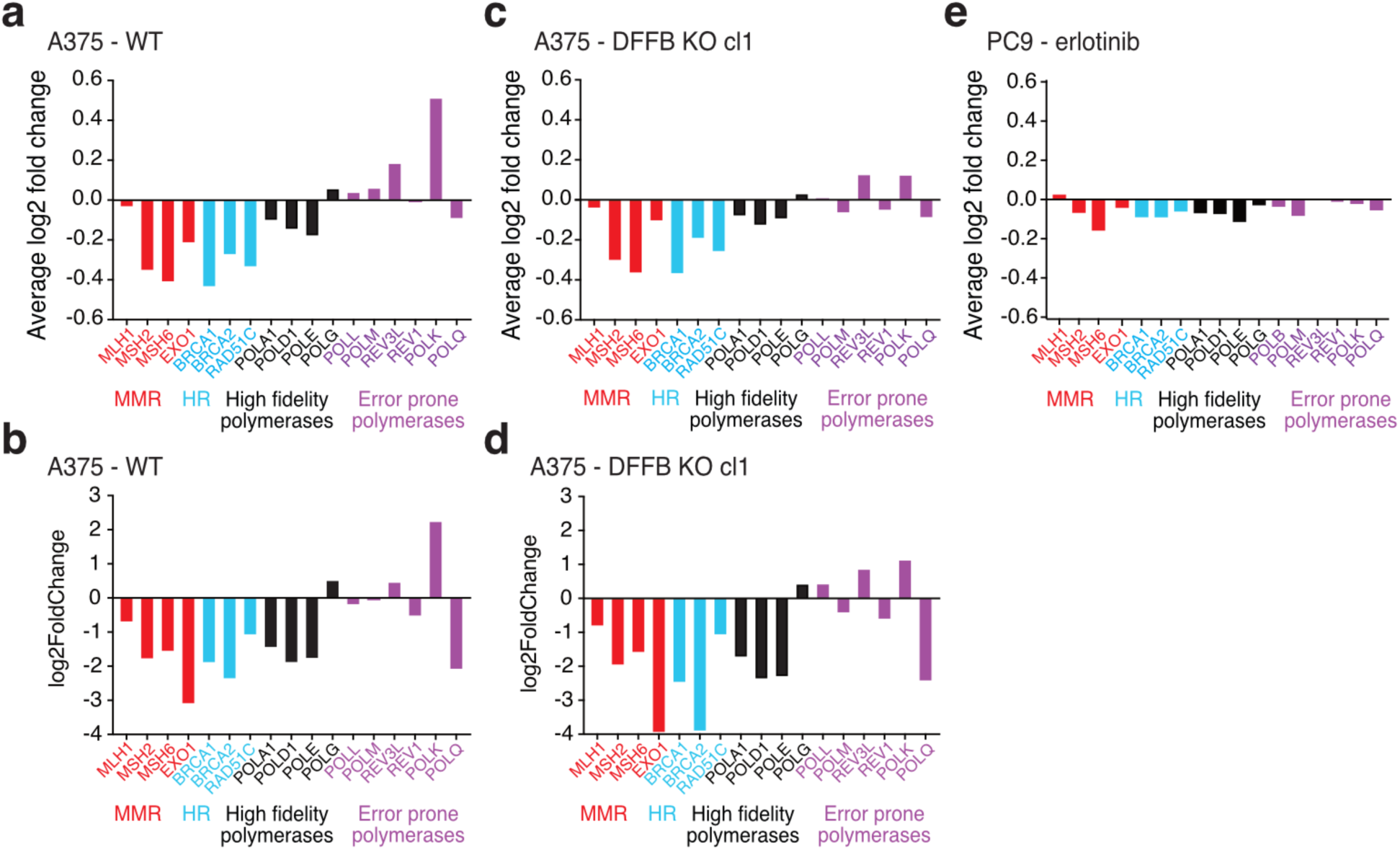
Stress-induced mutagenesis-associated transcriptional patterns. **a-e,** Comparison gene expression changes associated with stress-induced mutagenesis including downregulated repair genes, downregulated high fidelity polymerases, and upregulated error prone polymerases,^7^ in persister versus parental cells. **a-d**, A375 persister cells derived from 250 nM dabrafenib and 25 nM trametinib. **a,b,** A375 WT cells analyzed with (**a**) scRNAseq and (**b**) bulk RNAseq. **c,d,** A375 DFFB KO cells analyzed with (**c**) scRNAseq and (**d**) bulk RNAseq. cl, clone. **e,** PC9 persister cells treated with 2.5 μM erlotinib and analyzed with scRNAseq. Mismatch repair (MMR); homology-directed repair (HR).

**Extended Data Fig. 6.**
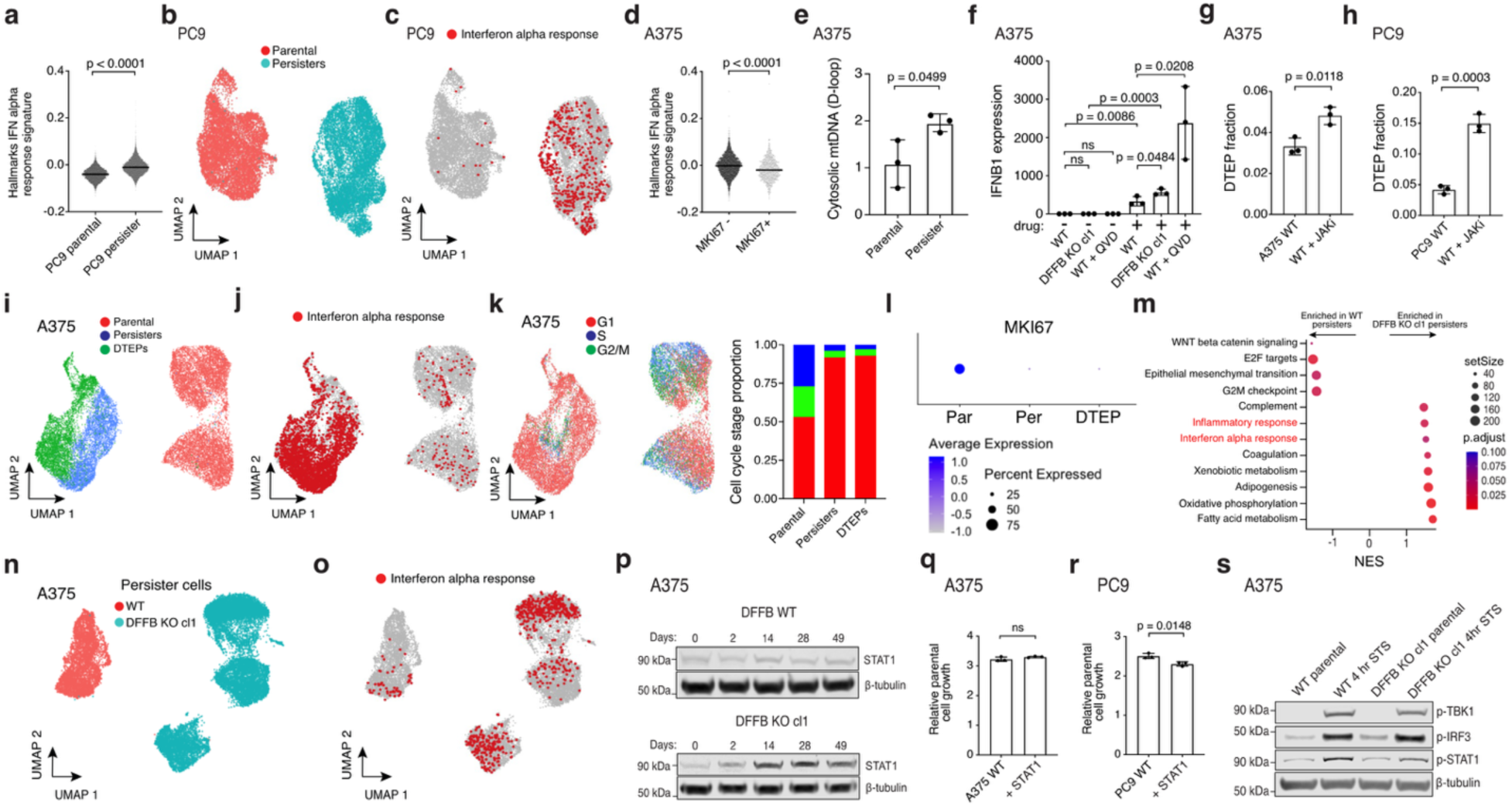
Increased STAT1 and ISGs restrict persister cell regrowth. **a**, ScRNAseq Hallmarks interferon alpha response gene set signature scores in PC9 cells. P values calculated with Mann-Whitney test. **b-c**, ScRNAseq UMAP visualizations of PC9 parental cells and persister cells derived from 2.5 μM erlotinib for 14 days (**b**) and cells with enriched Hallmarks interferon alpha response gene set highlighted in red (**c**). **d**, Hallmarks interferon alpha response gene set signature scores in A375 cycling (MKI67+) versus noncycling (MKI67-) DTEP cells. P value calculated with Mann-Whitney test. **e**, A375 cells analyzed for cytosolic mitochondrial DNA. **f**, IFNB1 mRNA expression in A375 cells treated with 1 μM dabrafenib and 100 nM trametinib for 6 days with and without 10 μM caspase inhibitor QVD treatment. cl, clone. **g**, DTEP formation from A375 WT persister cells treated with 250 nM dabrafenib and 25 nM trametinib for 2 weeks followed by the addition of 1 μM JAK inhibitor ruxolitinib for 5 weeks. **h**, DTEP formation from PC9 persister cells co-treated with 300 nM osimertinib and 5 μM ruxolitinib for 5 weeks. **i-k**, ScRNAseq UMAP visualizations of A375 parental cells, persister cells derived from treatment with 250 nM dabrafenib and 25 nM trametinib for 14 days, and DTEP cells treated for 9 weeks (**i**), cells with enriched Hallmarks interferon alpha response gene set highlighted in red (**j**), and cells colored by cell cycle stage (**k**). **l**, ScRNAseq MKI67 (Ki-67) expression in A375 parental (Par), persister (Per), and DTEP cells. **m**, Bulk RNAseq enriched Hallmarks gene sets between A375 DFFB KO versus WT persister cells. Normalized enrichment score, NES. **n-o**, scRNAseq UMAPs of A375 WT and DFFB KO persister cells (**n**) and cells with enriched Hallmarks interferon alpha response gene set highlighted in red (**o**)**. p**, A375 cells treated with 250 nM dabrafenib and 25 nM trametinib and assessed for STAT1 at multiple timepoints. **q-r**, A375 (**q**) and PC9 (**r**) parental cell proliferation with or without overexpressed STAT1. **s**, A375 cells treated with 1 μM staurosporine (STS) for 4 hours analyzed for activation of IFN signaling upstream of ISG expression. **e-h**,**q-r**, *n* = 3 biological replicates; mean ± s.d. is shown; P values calculated with two-tailed Student’s t-test. ns, not significant.

**Extended Data Fig. 7.**
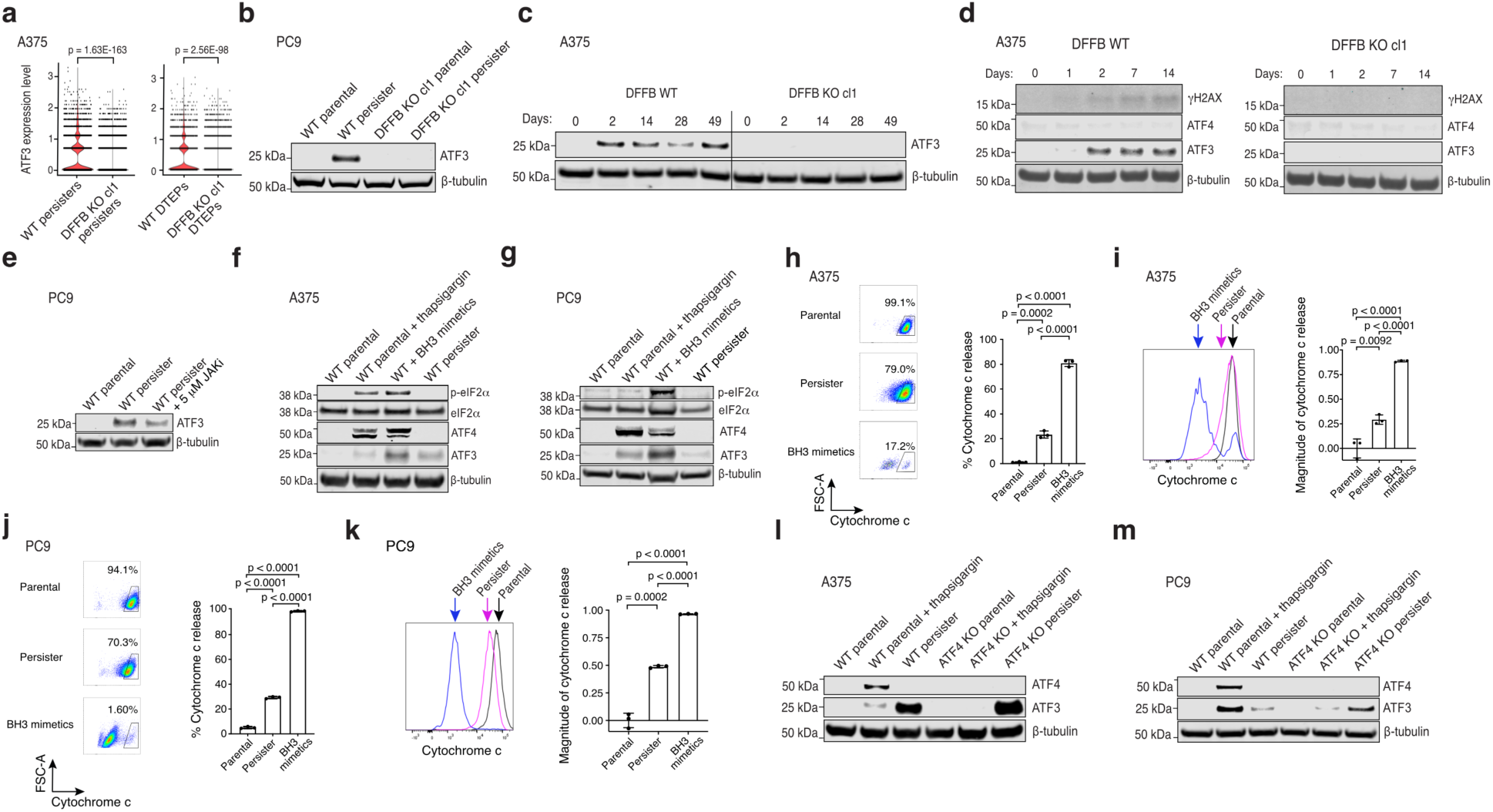
ATF3 is induced in persister cells independent of the integrated stress response. **a**, ATF3 expression levels between A375 WT and DFFB KO persister and DTEP cells measured with scRNAseq. P values calculated with the Wilcoxon Rank Sum test with Bonferroni correction. cl, clone. **b**, ATF3 expression in PC9 WT and DFFB KO cells treated with 500 nM osimertinib. **c**, A375 WT and DFFB KO cells treated with 250 nM dabrafenib and 25 nM trametinib analyzed for ATF3 at the indicated times. **d**, A375 WT and DFFB KO cells treated with 250 nM dabrafenib and 25 nM trametinib analyzed for γH2AX, ATF4, and ATF3 at the indicated times. **e**, PC9 persister cells derived from 500 nM osimertinib with or without 5 μM JAK inhibitor ruxolitinib and analyzed for ATF3 levels. **f**, A375 WT cells treated with ER stress inducer 1 μM thapsigargin for 4 hours, BH3 mimetics 5 μM ABT-737 and 10 μM S63845 for 2.5 hours followed by 24 hour recovery, and persister cells analyzed for ISR genes (phosphorylated (p) eIF2α, total eIF2α, ATF4) and ATF3. **g,** PC9 WT cells treated with 1 μM thapsigargin, BH3 mimetics 1.5 μM ABT-737 and 3 μM S63845 for 4 hours followed by 2 hour recovery, and persister cells analyzed for ISR genes and ATF3. **h-k,** Flow cytometry analysis of cytochrome C release in A375 (**h,i)** and PC9 (**j**,**k**) parental, persister, and BH3 mimetic-treated cells. **i** and **k,** Fractional loss of cytochrome C (geometric means) are plotted on the right graphs. *n* = 3 biological replicates; mean ± s.d. is shown; P values calculated with two-tailed Student’s t-test. **l,m,** A375 (**l**) and PC9 (**m**) WT and ATF4 CRISPR-depleted (pooled KO) persister cells were analyzed for ATF3 levels. For thapsigargin-induced ISR, ATF3 expression is decreased in ATF4-depleted cells. In persister cells (which lack the ISR) ATF3 is expressed despite ATF4-depletion.

**Extended Data Fig. 8.**
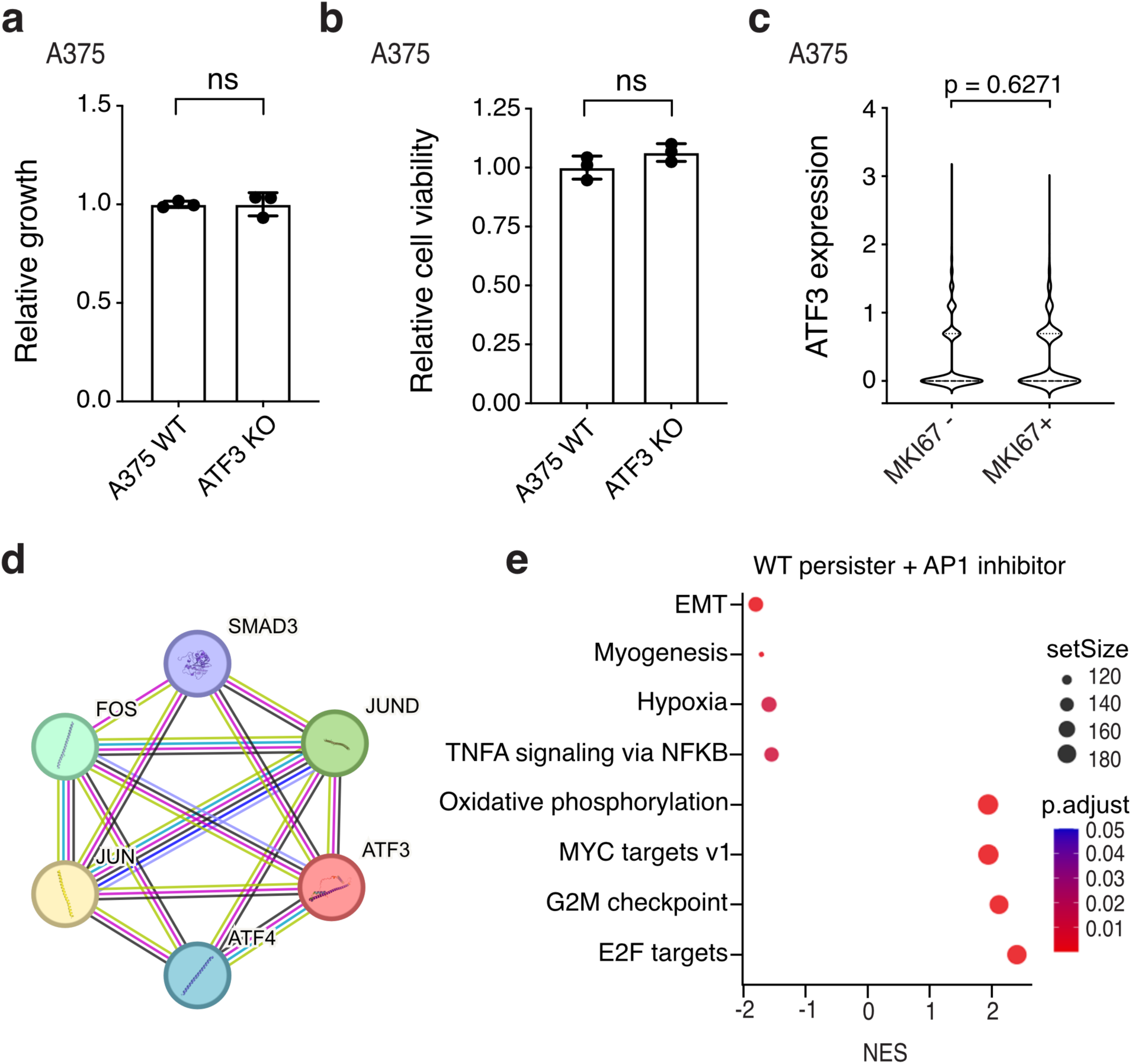
Additional ATF3 and AP1 data. **a**,**b**, A375 ATF3-depleted (pooled KO) cells assessed for parental cell proliferation (**a**) and persister cell formation (**b**). **c**, A375 WT DTEP cycling (MKI67+) and noncycling (MKI67-) cell ATF3 scRNAseq expression. P values calculated with the Wilcoxon Rank Sum test. **d**, STRING analysis of ATF3 interactions. **e**, Bulk RNAseq gene set enrichment analysis of A375 WT persister cells versus persister cells co-treated with 20 μM AP1 inhibitor T-5224. *n* = 3 biological replicates. **a**,**b**, *n* = 3 biological replicates; mean ± s.d. is shown; P values calculated with two-tailed Student’s t-test. ns, not significant.

**Extended Data Fig. 9.**
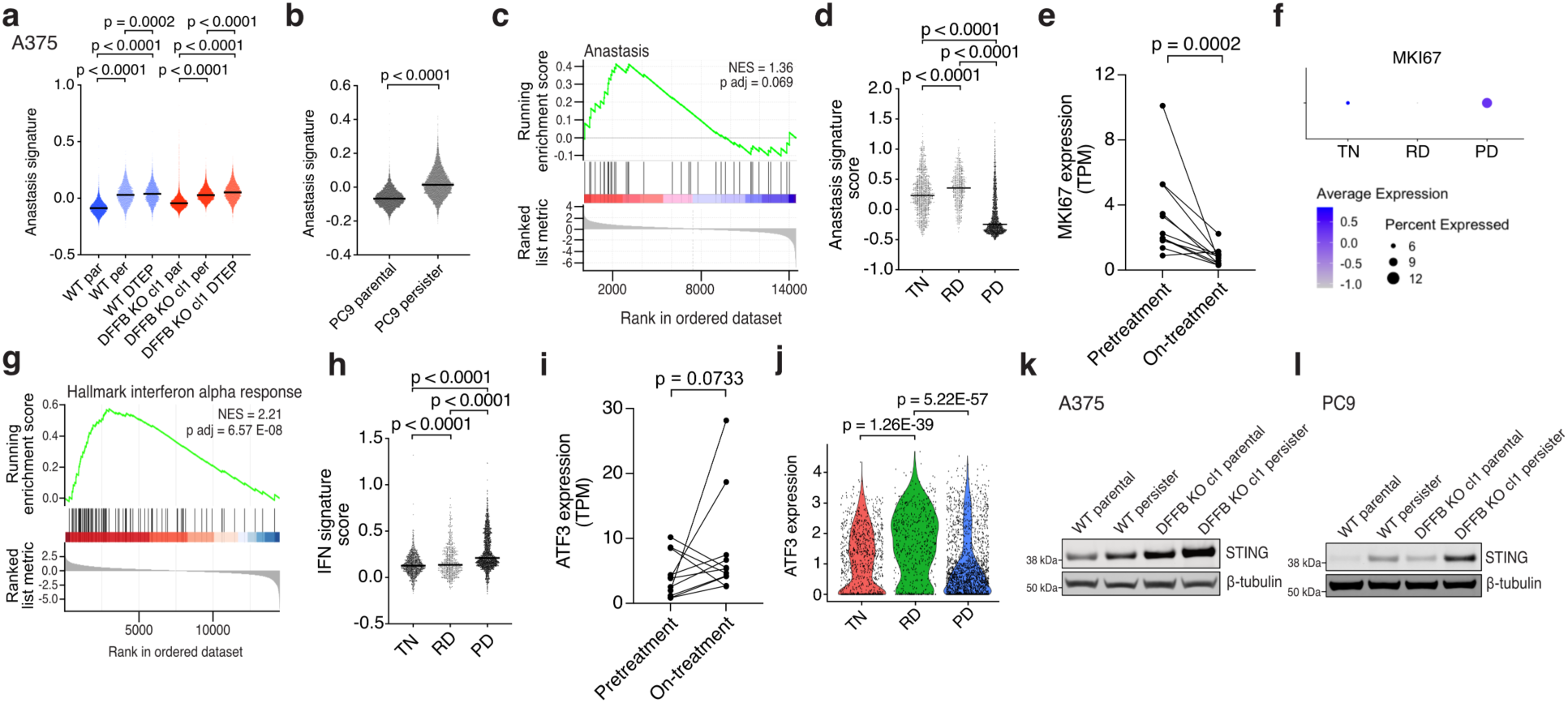
Comparison of persister and DTEP cell signatures with targeted therapy-treated patient tumours. **a**,**b**, ScRNAseq anastasis gene set signature scores in A375 parental, persister, and DTEP timepoint DFFB WT and KO cells (**a**) and PC9 parental and persister cells (**b**). P values calculated with Mann-Whitney test. Anastasis remains elevated at the DTEP timepoint, indicating that DTEP cells remain drug stressed. cl, clone. **c**, Anastasis gene set expression analyzed in on-treatment versus pretreatment patient melanoma tumours treated with BRAF +/- MEK targeted therapy. Normalized enrichment score, NES. P value calculated with a permutation test with family-wise error rate correction. **d**, Anastasis gene set signature scores in treatment naïve (TN), residual disease (RD), and progressive disease (PD) patient non-small cell lung cancer tumours treated with EGFR targeted therapy. **e**, MKI67 (Ki-67) expression in pretreatment and on-treatment melanoma patient tumours. **f**, Ki-67 expression in each lung cancer treatment response stage. **g-h**, Hallmarks interferon alpha response gene set expression analyzed in on-treatment versus pretreatment patient melanoma (**g**) and lung cancer tumour treatment response stages (**h**). **i**, ATF3 expression in pretreatment and on-treatment patient melanoma tumours. **j**, ATF3 expression in each lung cancer treatment response stage. P value calculated with the Wilcoxon Rank Sum test with Bonferroni correction. **k,l**, A375 (**k**) and PC9 (**l**) WT and DFFB KO parental and persister cells analyzed for STING expression. **a,b,d**,**h**, P values calculated with Mann-Whitney test. **e**,**i**, *n* = 11 patients. P values calculated with paired ratio t-test.

## References

1. Dagogo-Jack, I. & Shaw, A. T. Tumour heterogeneity and resistance to cancer therapies. Nat. Rev. Clin. Oncol. 15, 81–94 (2018).

2. Romano, E. et al. Identification of Multiple Mechanisms of Resistance to Vemurafenib in a Patient with BRAFV600E-Mutated Cutaneous Melanoma Successfully Rechallenged after Progression. Clin. Cancer Res. 19, 5749–5757 (2013).

3. Juric, D. et al. Convergent loss of PTEN leads to clinical resistance to a PI(3)Kα inhibitor. Nature 518, 240–244 (2015).

4. Mullard, A. Stemming the tide of drug resistance in cancer. Nat. Rev. Drug Discov. 19, 221– 223 (2020).

5. Hata, A. N. et al. Tumor cells can follow distinct evolutionary paths to become resistant to epidermal growth factor receptor inhibition. Nat. Med. 22, 262–269 (2016).

6. Ramirez, M. et al. Diverse drug-resistance mechanisms can emerge from drug-tolerant cancer persister cells. Nat. Commun. 7, 10690 (2016).

7. Russo, M. et al. Adaptive mutability of colorectal cancers in response to targeted therapies. Science 366, 1473–1480 (2019).

8. Cipponi, A. et al. MTOR signaling orchestrates stress-induced mutagenesis, facilitating adaptive evolution in cancer. Science 368, 1127–1131 (2020).

9. Channathodiyil, P., et al. Escape from G1 arrest during acute MEK inhibition drives the acquisition of drug resistance. NAR Cancer 4, zcac032 (2022).

10. Sharma, S. V. et al. A Chromatin-Mediated Reversible Drug-Tolerant State in Cancer Cell Subpopulations. Cell 141, 69–80 (2010).

11. Hangauer, M. J. et al. Drug-tolerant persister cancer cells are vulnerable to GPX4 inhibition. Nature 551, 247–250 (2017).

12. Tang, H. L. et al. Cell survival, DNA damage, and oncogenic transformation after a transient and reversible apoptotic response. Mol. Biol. Cell 23, 2240–2252 (2012).

13. Kalkavan, H. et al. Sublethal cytochrome c release generates drug-tolerant persister cells. Cell 185, 3356–3374.e22 (2022).

14. Berthenet, K. et al. Failed Apoptosis Enhances Melanoma Cancer Cell Aggressiveness. Cell Rep. 31, 107731 (2020).

15. Nano, M., Mondo, J. A., Harwood, J., Balasanyan, V. & Montell, D. J. Cell survival following direct executioner-caspase activation. Proc. Natl. Acad. Sci. 120, e2216531120 (2023).

16. Liu, X. et al. Self-inflicted DNA double-strand breaks sustain tumorigenicity and stemness of cancer cells. Cell Res. 27, 764–783 (2017).

17. Lovric, M. M. & Hawkins, C. J. TRAIL treatment provokes mutations in surviving cells. Oncogene 29, 5048–5060 (2010).

18. Ichim, G. et al. Limited Mitochondrial Permeabilization Causes DNA Damage and Genomic Instability in the Absence of Cell Death. Mol. Cell 57, 860–872 (2015).

19. Ali, M. et al. Small-molecule targeted therapies induce dependence on DNA double-strand break repair in residual tumor cells. Sci. Transl. Med. 14, eabc7480 (2022).

20. Benada, J., Alsowaida, D., Megeney, L. A. & Sørensen, C. S. Self-inflicted DNA breaks in cell differentiation and cancer. Trends Cell Biol. 33, 850–859 (2023).

21. Iglesias-Guimarais, V. et al. Chromatin Collapse during Caspase-dependent Apoptotic Cell Death Requires DNA Fragmentation Factor, 40-kDa Subunit-/Caspase-activated Deoxyribonuclease-mediated 3ʹ-OH Single-strand DNA Breaks*. J. Biol. Chem. 288, 9200–9215 (2013).

22. Larsen, B. D. & Sørensen, C. S. The caspase-activated DNase: apoptosis and beyond. FEBS J. 284, 1160–1170 (2017).

23. Liu, X., Zou, H., Widlak, P., Garrard, W. & Wang, X. Activation of the Apoptotic Endonuclease DFF40 (Caspase-activated DNase or Nuclease): OLIGOMERIZATION AND DIRECT INTERACTION WITH HISTONE H1*. J. Biol. Chem. 274, 13836–13840 (1999).

24. Haimovici, A. et al. Spontaneous activity of the mitochondrial apoptosis pathway drives chromosomal defects, the appearance of micronuclei and cancer metastasis through the Caspase-Activated DNAse. Cell Death Dis. 13, 315 (2022).

25. Kula, T. et al. T-Scan: A Genome-wide Method for the Systematic Discovery of T Cell Epitopes. Cell 178, 1016–1028.e13 (2019).

26. Meiss, G., Scholz, S. R., Korn, C., Gimadutdinow, O. & Pingoud, A. Identification of functionally relevant histidine residues in the apoptotic nuclease CAD. Nucleic Acids Res. 29, 3901–3909 (2001).

27. Miles, M. A. & Hawkins, C. J. Executioner caspases and CAD are essential for mutagenesis induced by TRAIL or vincristine. Cell Death Dis. 8, e3062–e3062 (2017).

28. Miles, M. A., Harris, M. A. & Hawkins, C. J. Proteasome inhibitors trigger mutations via activation of caspases and CAD, but mutagenesis provoked by the HDAC inhibitors vorinostat and romidepsin is caspase/CAD-independent. Apoptosis 24, 404–413 (2019).

29. Isozaki, H. et al. Therapy-induced APOBEC3A drives evolution of persistent cancer cells. Nature 620, 393–401 (2023).

30. Ivashkiv, L. B. & Donlin, L. T. Regulation of type I interferon responses. Nat. Rev. Immunol. 14, 36–49 (2014).

31. Hu, J. et al. STING inhibits the reactivation of dormant metastasis in lung adenocarcinoma. Nature 616, 806–813 (2023).

32. Killarney, S. T. et al. Executioner caspases restrict mitochondrial RNA-driven Type I IFN induction during chemotherapy-induced apoptosis. Nat. Commun. 14, 1399 (2023).

33. Cheon, H., Wang, Y., Wightman, S. M., Jackson, M. W. & Stark, G. R. How cancer cells make and respond to interferon-I. Trends Cancer 9, 83–92 (2023).

34. Cheon, H. & Stark, G. R. Unphosphorylated STAT1 prolongs the expression of interferon-induced immune regulatory genes. Proc. Natl. Acad. Sci. 106, 9373–9378 (2009).

35. Moeed, A. et al. The Caspase-Activated DNase drives inflammation and contributes to defense against viral infection. Cell Death Differ. 1–14 (2024) doi:10.1038/s41418-024-01320-7.

36. Rongvaux, A. et al. Apoptotic Caspases Prevent the Induction of Type I Interferons by Mitochondrial DNA. Cell 159, 1563–1577 (2014).

37. White, M. J. et al. Apoptotic Caspases Suppress mtDNA-Induced STING-Mediated Type I IFN Production. Cell 159, 1549–1562 (2014).

38. Larsen, B. D. et al. Cancer cells use self-inflicted DNA breaks to evade growth limits imposed by genotoxic stress. Science 376, 476–483 (2022).

39. Larsen, B. D. et al. Caspase 3/caspase-activated DNase promote cell differentiation by inducing DNA strand breaks. Proc. Natl. Acad. Sci. 107, 4230–4235 (2010).

40. Labzin, L. I. et al. ATF3 Is a Key Regulator of Macrophage IFN Responses. J. Immunol. 195, 4446–4455 (2015).

41. Cai, X., Xu, Y., Kim, Y.-M., Loureiro, J. & Huang, Q. PIKfyve, a Class III Lipid Kinase, Is Required for TLR-Induced Type I IFN Production via Modulation of ATF3. J. Immunol. 192, 3383–3389 (2014).

42. Badu, P., Baniulyte, G., Sammons, M. A. & Pager, C. T. Activation of ATF3 via the integrated stress response pathway regulates innate immune response to restrict Zika virus. J. Virol. 98, e01055–24 (2024).

43. Mao, J. et al. Reactivation of senescence-associated endogenous retroviruses by ATF3 drives interferon signaling in aging. Nat. Aging 4, 1794–1812 (2024).

44. Amundson, S. A. et al. Fluorescent cDNA microarray hybridization reveals complexity and heterogeneity of cellular genotoxic stress responses. Oncogene 18, 3666–3672 (1999).

45. Ho, H. H., Antoniv, T. T., Ji, J.-D. & Ivashkiv, L. B. Lipopolysaccharide-Induced Expression of Matrix Metalloproteinases in Human Monocytes Is Suppressed by IFN-γ via Superinduction of ATF-3 and Suppression of AP-11. J. Immunol. 181, 5089–5097 (2008).

46. Jiang, H.-Y. et al. Activating Transcription Factor 3 Is Integral to the Eukaryotic Initiation Factor 2 Kinase Stress Response. Mol. Cell. Biol. 24, 1365–1377 (2004).

47. Le, T. T. et al. Etoposide promotes DNA loop trapping and barrier formation by topoisomerase II. Nat. Chem. Biol. 19, 641–650 (2023).

48. Costa-Mattioli, M. & Walter, P. The integrated stress response: From mechanism to disease. Science 368, eaat5314 (2020).

49. Ku, H.-C. & Cheng, C.-F. Master Regulator Activating Transcription Factor 3 (ATF3) in Metabolic Homeostasis and Cancer. Front. Endocrinol. 11, (2020).

50. Larsen, S. B. et al. Establishment, maintenance, and recall of inflammatory memory. Cell Stem Cell 28, 1758–1774.e8 (2021).

51. Kwong, L. N. et al. Co-clinical assessment identifies patterns of BRAF inhibitor resistance in melanoma. J. Clin. Invest. 125, 1459–1470 (2015).

52. Maynard, A. et al. Therapy-Induced Evolution of Human Lung Cancer Revealed by Single-Cell RNA Sequencing. Cell 182, 1232–1251.e22 (2020).

53. Russo, M. et al. Cancer drug-tolerant persister cells: from biological questions to clinical opportunities. Nat. Rev. Cancer 24, 694–717 (2024).

54. Hu, Q. et al. Elevated cleaved caspase-3 is associated with shortened overall survival in several cancer types. Int. J. Clin. Exp. Pathol. 7, 5057–5070 (2014).

55. Yang, C., Tian, C., Hoffman, T. E., Jacobsen, N. K. & Spencer, S. L. Melanoma subpopulations that rapidly escape MAPK pathway inhibition incur DNA damage and rely on stress signalling. Nat. Commun. 12, 1747 (2021).

56. Zhang, J. et al. Resistance to DNA fragmentation and chromatin condensation in mice lacking the DNA fragmentation factor 45. Proc. Natl. Acad. Sci. 95, 12480–12485 (1998).

57. Kawane, K. et al. Impaired thymic development in mouse embryos deficient in apoptotic DNA degradation. Nat. Immunol. 4, 138–144 (2003).

58. Woo, E.-J. et al. Structural Mechanism for Inactivation and Activation of CAD/DFF40 in the Apoptotic Pathway. Mol. Cell 14, 531–539 (2004).

59. Samejima, K. & Earnshaw, W. C. Trashing the genome: the role of nucleases during apoptosis. Nat. Rev. Mol. Cell Biol. 6, 677–688 (2005).

60. Ma, F. et al. Positive feedback regulation of type I interferon by the interferon-stimulated gene STING. EMBO Rep. 16, 202–212 (2015).

## Methods References

61. Zheng, G. X. Y. et al. Massively parallel digital transcriptional profiling of single cells. Nat. Commun. 8, 14049 (2017).

62. Satija, R., Farrell, J. A., Gennert, D., Schier, A. F. & Regev, A. Spatial reconstruction of single-cell gene expression data. Nat. Biotechnol. 33, 495–502 (2015).

63. Butler, A., Hoffman, P., Smibert, P., Papalexi, E. & Satija, R. Integrating single-cell transcriptomic data across different conditions, technologies, and species. Nat. Biotechnol. 36, 411–420 (2018).

64. Stuart, T. et al. Comprehensive Integration of Single-Cell Data. Cell 177, 1888–1902.e21 (2019).

65. Kowalczyk, M. S. et al. Single-cell RNA-seq reveals changes in cell cycle and differentiation programs upon aging of hematopoietic stem cells. Genome Res. 25, 1860–1872 (2015).

66. Yu, G., Wang, L.-G., Han, Y. & He, Q.-Y. clusterProfiler: an R Package for Comparing Biological Themes Among Gene Clusters. OMICS J. Integr. Biol. 16, 284–287 (2012).

67. Aibar, S. et al. SCENIC: single-cell regulatory network inference and clustering. Nat. Methods 14, 1083–1086 (2017).

68. Villalobos-Ortiz, M., Ryan, J., Mashaka, T. N., Opferman, J. T. & Letai, A. BH3 profiling discriminates on-target small molecule BH3 mimetics from putative mimetics. Cell Death Differ. 27, 999–1007 (2020).

69. McKenna, A. et al. The Genome Analysis Toolkit: A MapReduce framework for analyzing next-generation DNA sequencing data. Genome Res. 20, 1297–1303 (2010).

70. DePristo, M. A. et al. A framework for variation discovery and genotyping using next-generation DNA sequencing data. Nat. Genet. 43, 491–498 (2011).

71. Auwera, G. A. V. der & O’Connor, B. D. Genomics in the Cloud: Using Docker, GATK, and WDL in Terra. (O’Reilly Media, Inc., 2020).

72. Li, H. & Durbin, R. Fast and accurate short read alignment with Burrows–Wheeler transform. Bioinformatics 25, 1754–1760 (2009).

73. Mayakonda, A., Lin, D.-C., Assenov, Y., Plass, C. & Koeffler, H. P. Maftools: efficient and comprehensive analysis of somatic variants in cancer. Genome Res. 28, 1747–1756 (2018).

74. Victorelli, S. et al. Apoptotic stress causes mtDNA release during senescence and drives the SASP. Nature 622, 627–636 (2023).

75. Kim, E. & Hart, T. Improved analysis of CRISPR fitness screens and reduced off-target effects with the BAGEL2 gene essentiality classifier. Genome Med. 13, 2 (2021).

