## Supplementary information for "DFFB suppresses interferon to enable cancer persister cell regrowth"

**Supplementary Figure 1: Unprocessed western blot images.**

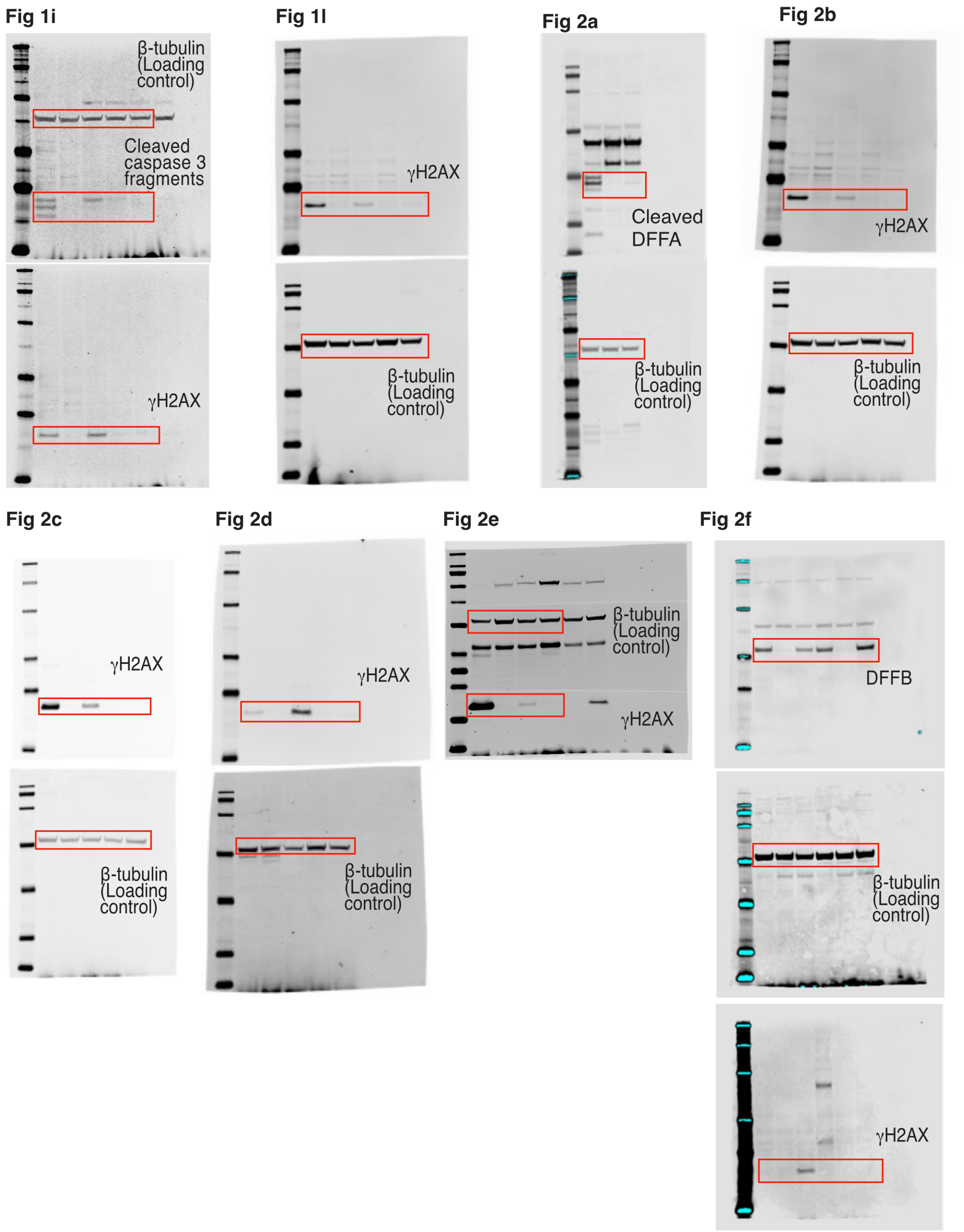

**Fig 3f**

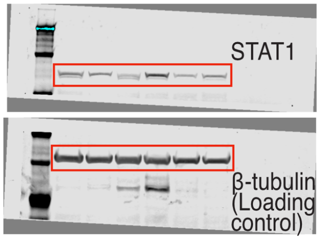


**Fig 3g**

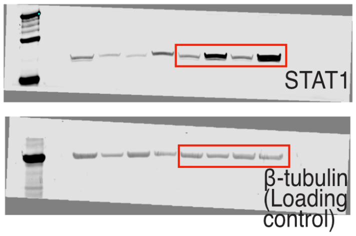


**Fig 3h**

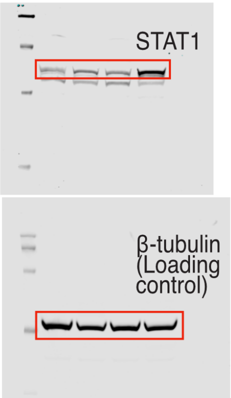


**Fig 3i**

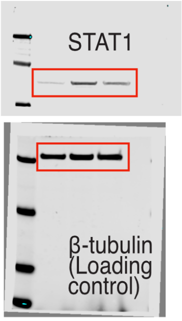


**Fig 3l**

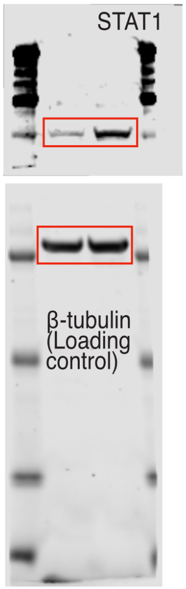


**Fig 3j**

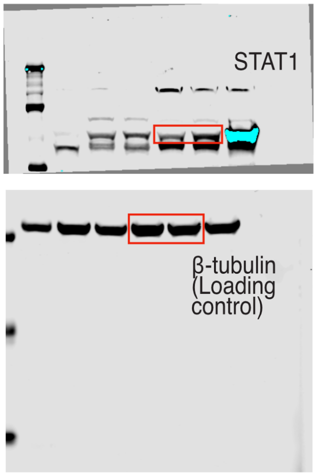


**Fig 4a**

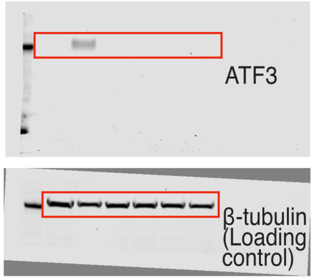


**Fig 4b**

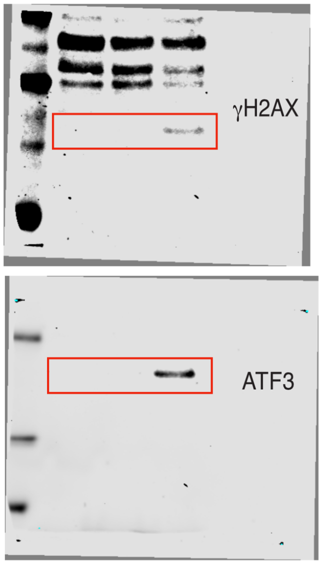


**Fig 4b (continued)**

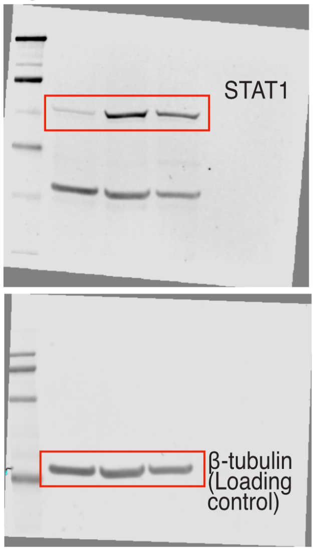


**Fig 4c**

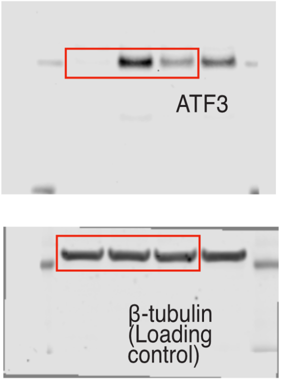


**Fig 4d**

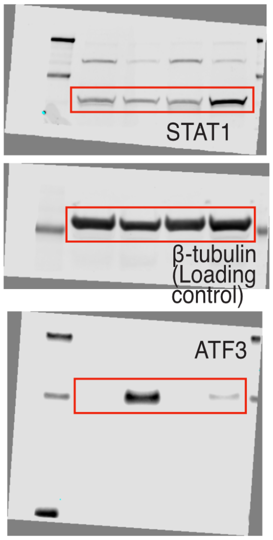

Fig 4f

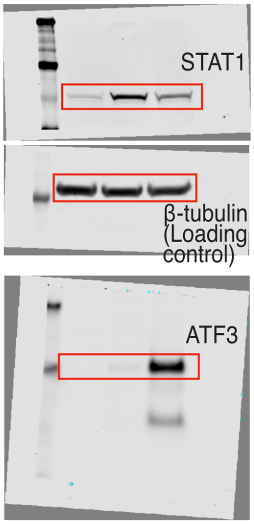


Extended Data Fig 2c

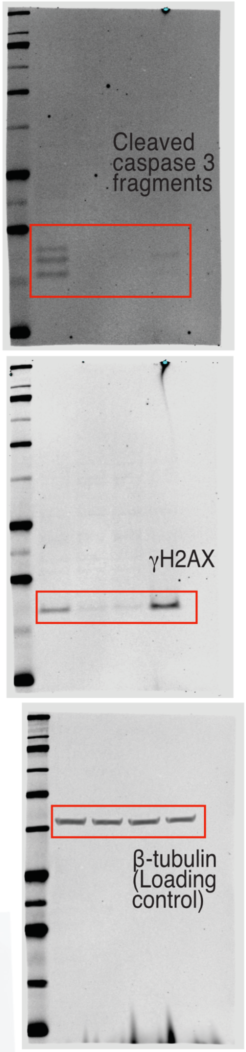


Extended Data Fig 2d

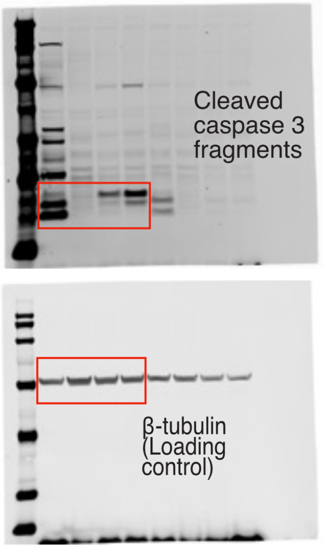


Extended Data Fig 2e

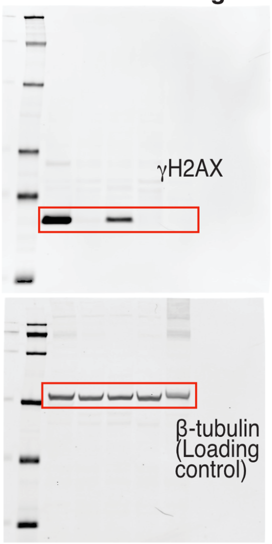


Extended Data Fig 2f

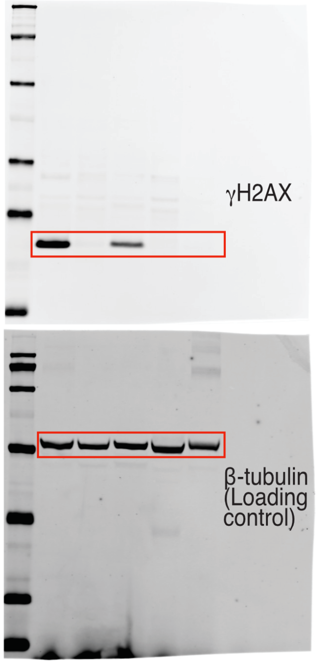


Extended Data Fig 2g

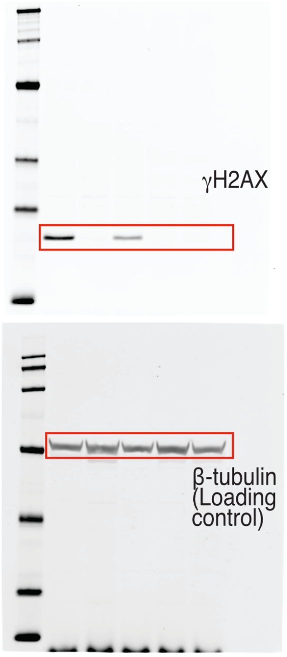

**Extended Data Fig 2i**

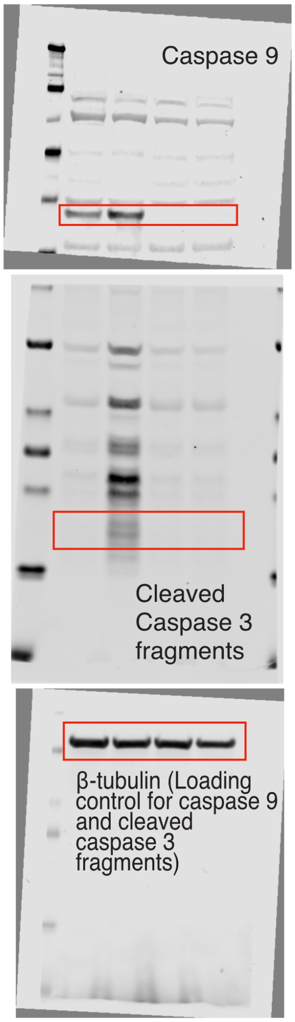


**Extended Data Fig 2i (continued)**

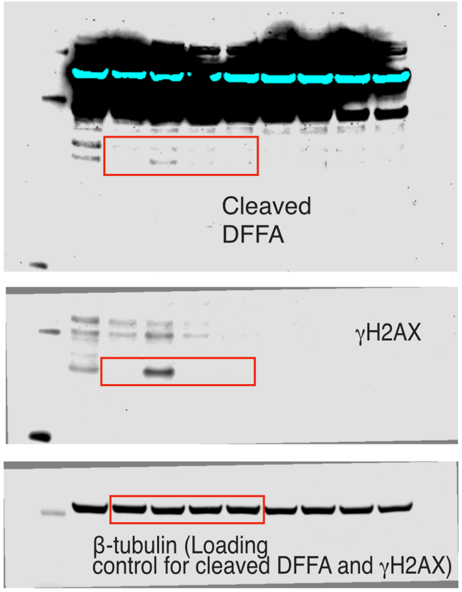


**Extended Data Fig 2j**

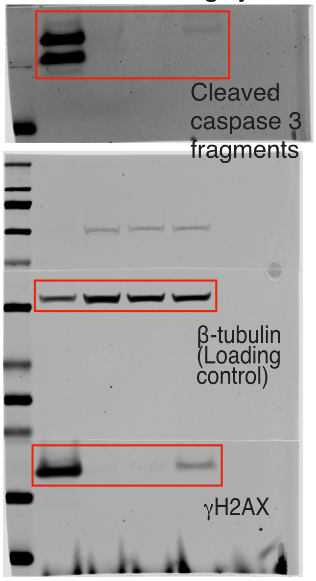


**Extended Data Fig 2k**

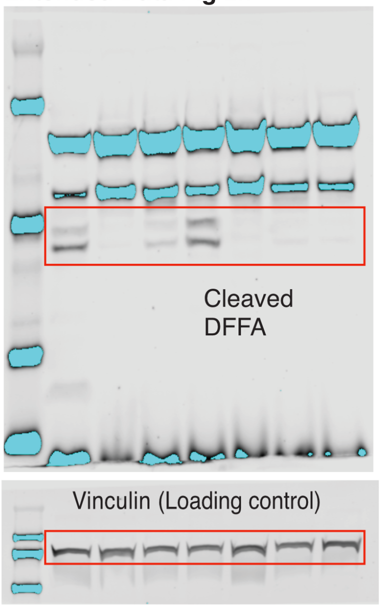


**Extended Data Fig 3a**

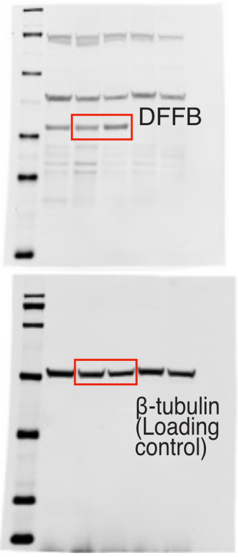


**Extended Data Fig 3b**

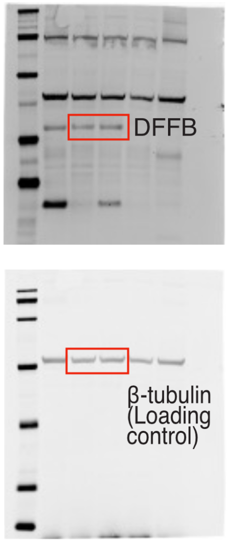


**Extended Data Fig 3c**

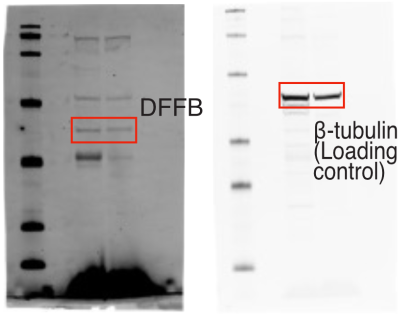

**Extended Data Fig 3d**

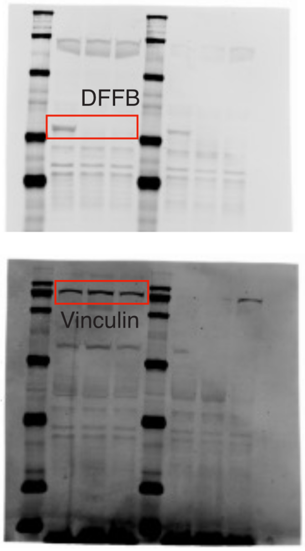


**Extended Data Fig 3e**

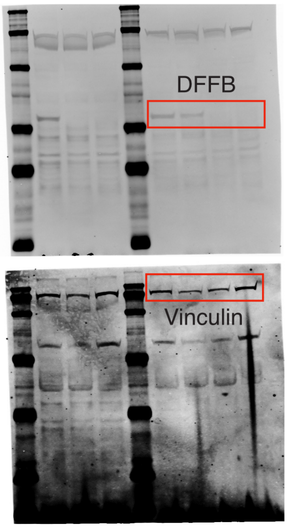


**Extended Data Fig 3f**

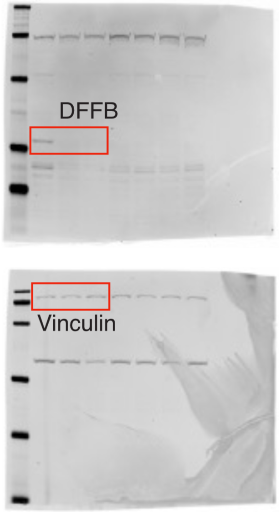


**Extended Data Fig 3g**

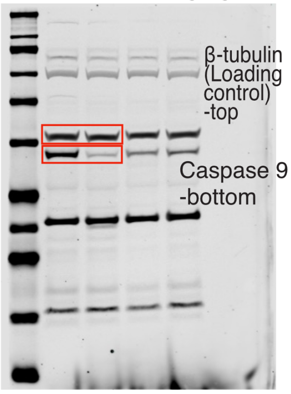


**Extended Data Fig 3h**

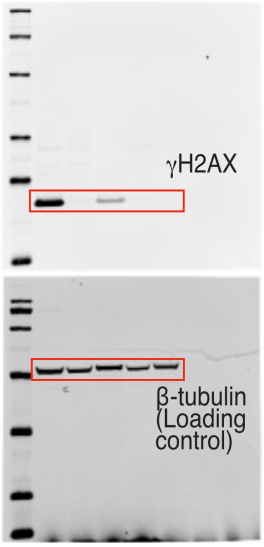


**Extended Data Fig 4q**

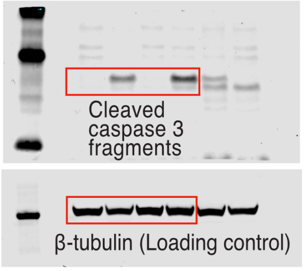


**Extended Data Fig 6o**

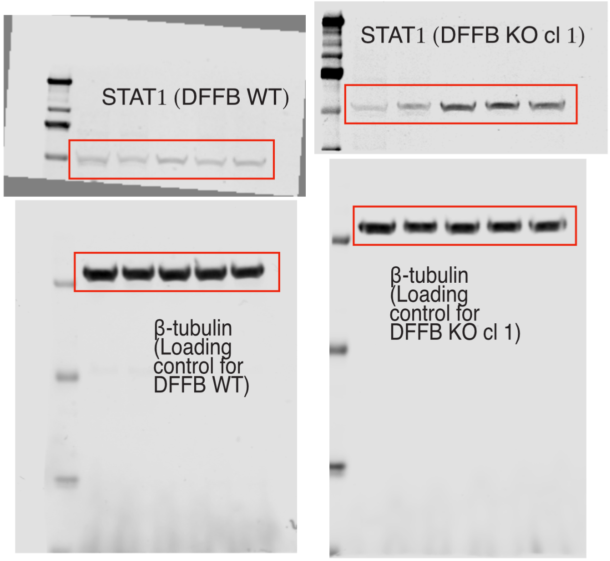

**Extended Data Fig 6r**

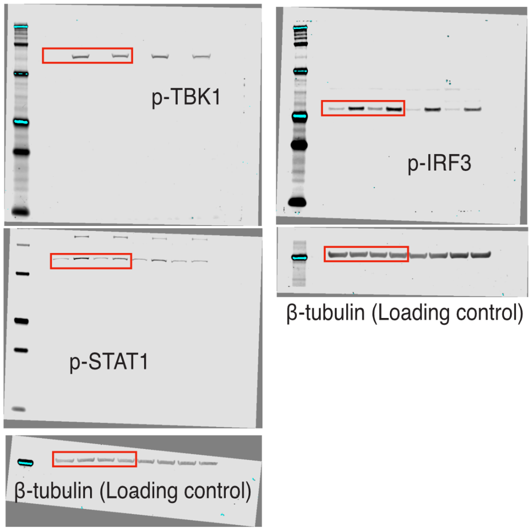


**Extended Data Fig 7b**

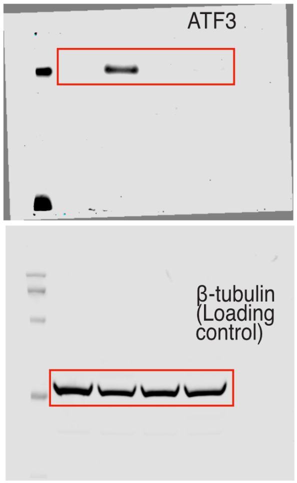


**Extended Data Fig 7c**

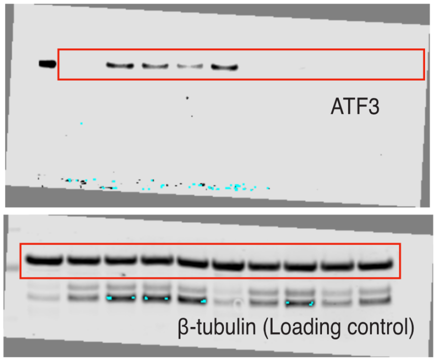


**Extended Data Fig 7d**

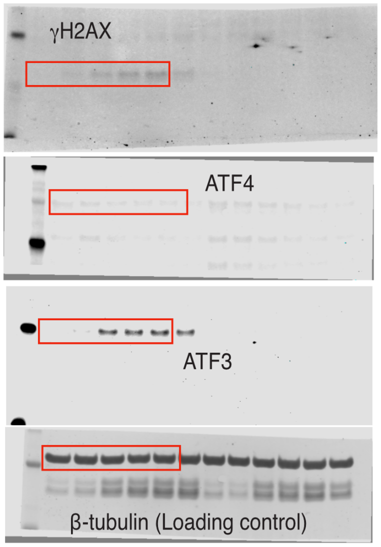


**Extended Data Fig 7d (continued)**

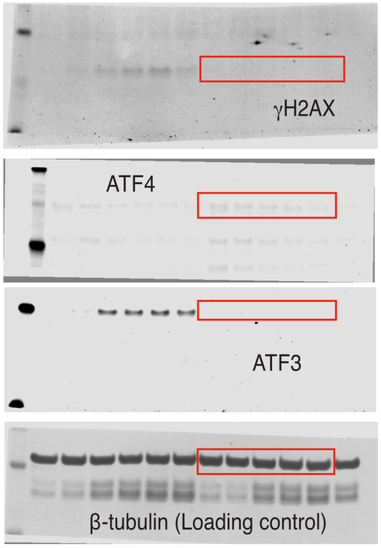

**Extended Data Fig 7e**

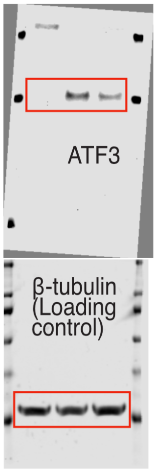


**Extended Data Fig 7f**

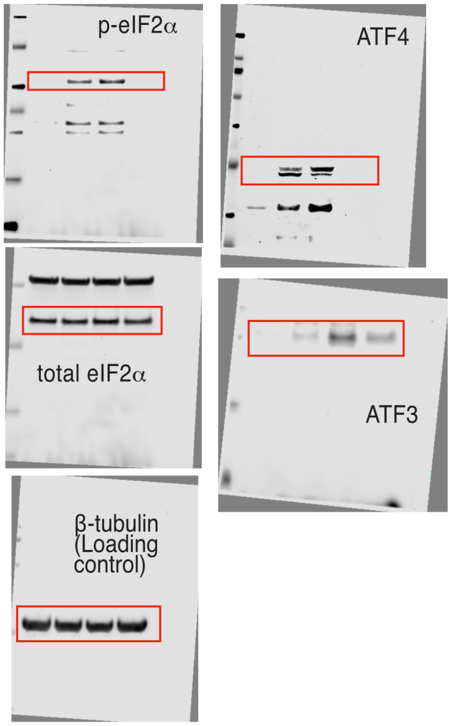


**Extended Data Fig 7g**

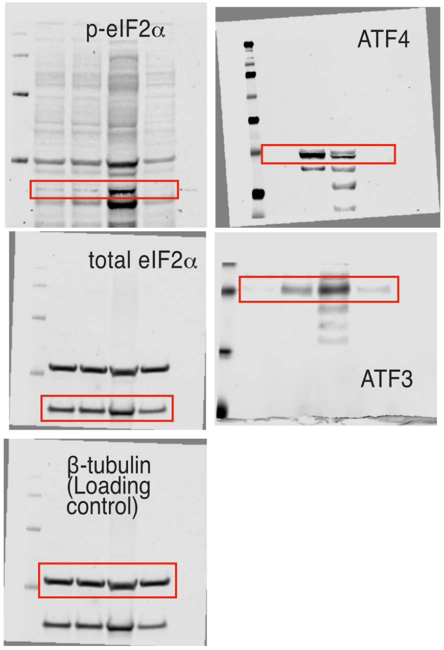


**Extended Data Fig 7l**

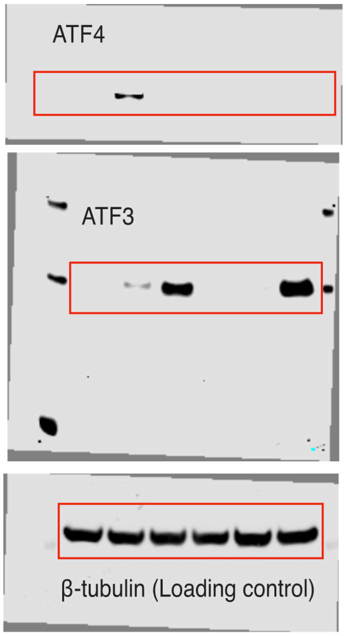


**Extended Data Fig 7m**

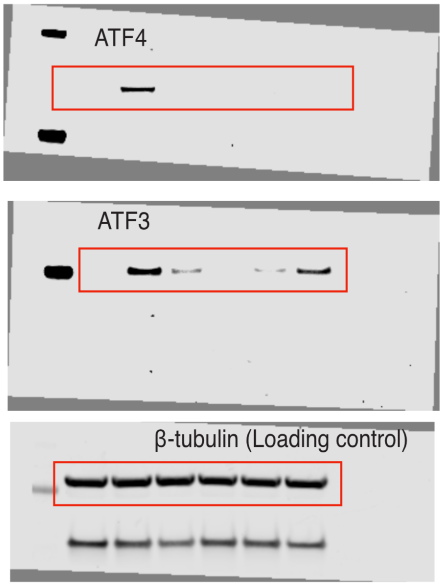


**Extended Data Fig 9k**

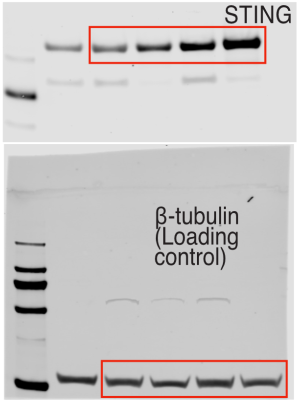


**Extended Data Fig 9l**

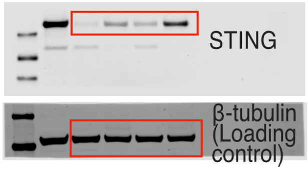

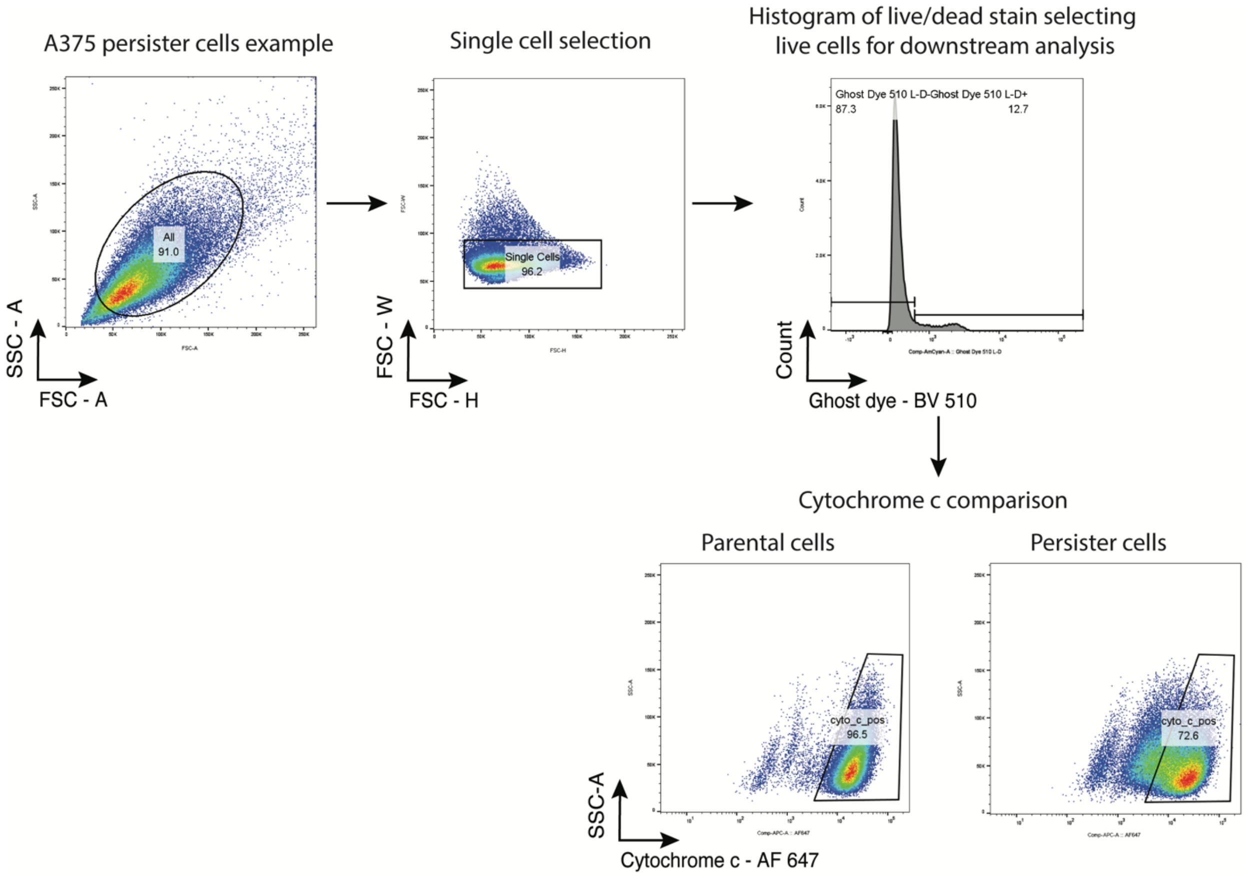


**Supplementary Figure 2: Mitochondrial release of cytochrome c flow cytometry assay schematic.** Gating strategy used for A375 parental cells and persister cells derived from treatment with 250 nM dabrafenib and 25 nM trametinib in Figure 1g. See methods for details of this assay.

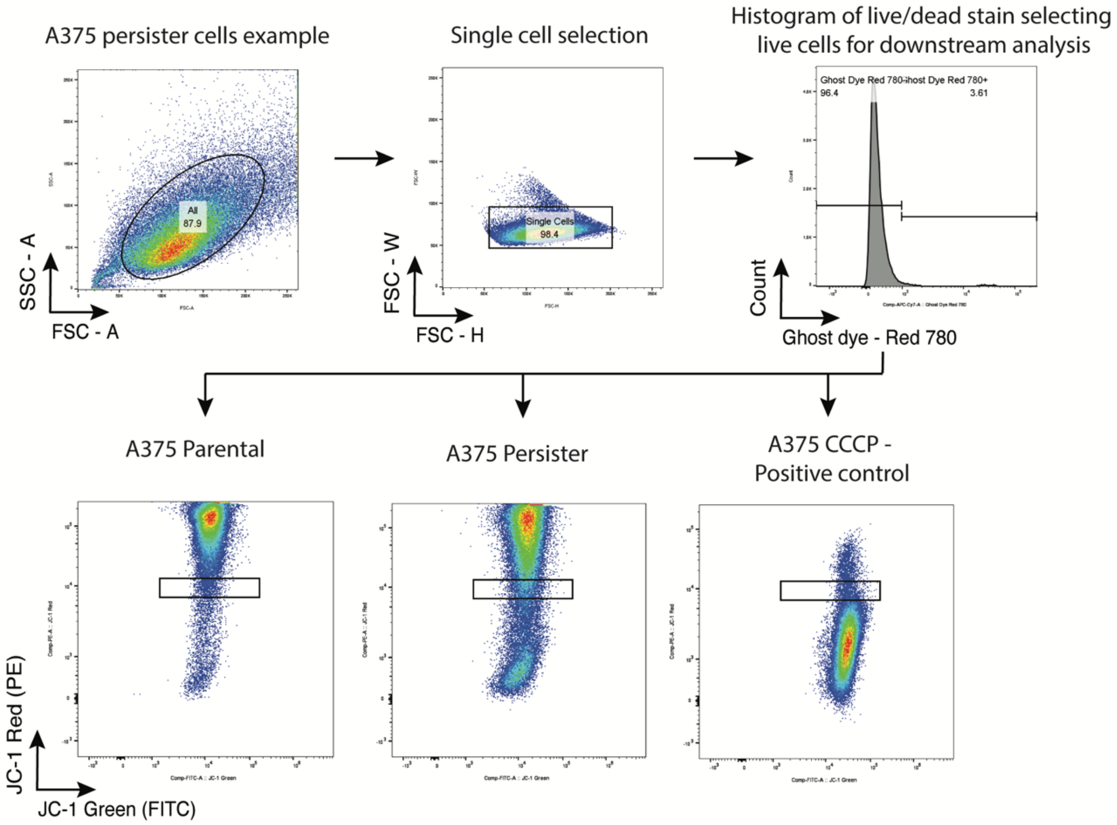


**Supplementary Figure 3: Flow cytometry schematic for JC-1 indicator dye-based assessment of loss of mitochondrial potential.** Gating strategy used for Extended Data Figure 3a. Representative flow cytometry gating for loss of mitochondrial potential in A375 parental cells and persister cells treated with 250 nM dabrafenib and 25 nM trametinib. Results from triplicate biological replicates are presented in Fig. 1h. For the positive control, cells were treated with mitochondrial oxidative phosphorylation uncoupler carbonyl cyanide m-chlorophenylhydrazone (CCCP) (50 μM) for 5 minutes. Gating was set to compare relative levels of incomplete loss of mitochondrial potential, indicated by intermediate levels of loss of JC-1 red fluorescence, as illustrated.

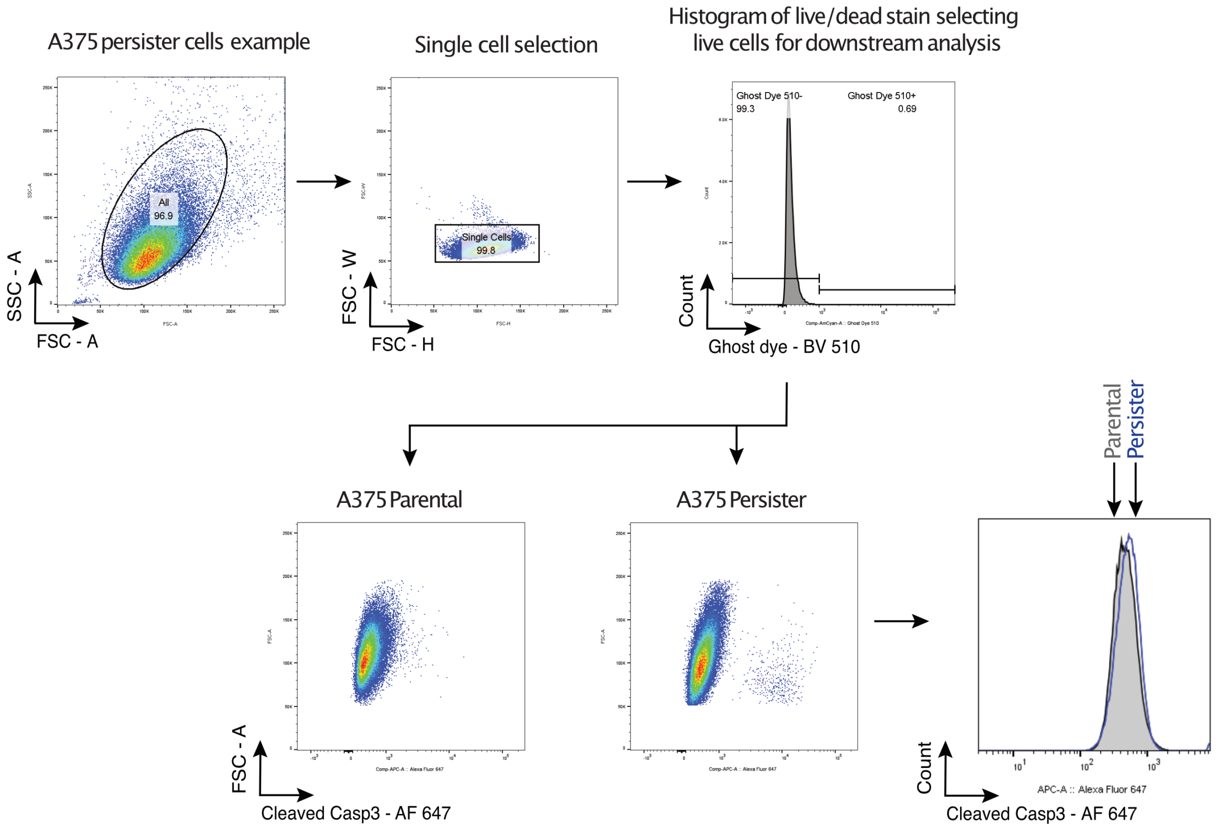


**Supplementary Figure 4: Cleaved caspase 3 flow cytometry schematic.** Gating strategy used for A375 parental cells and persister cells derived from treatment with 250 nM dabrafenib and 25 nM trametinib in Extended Data Figure 2a.

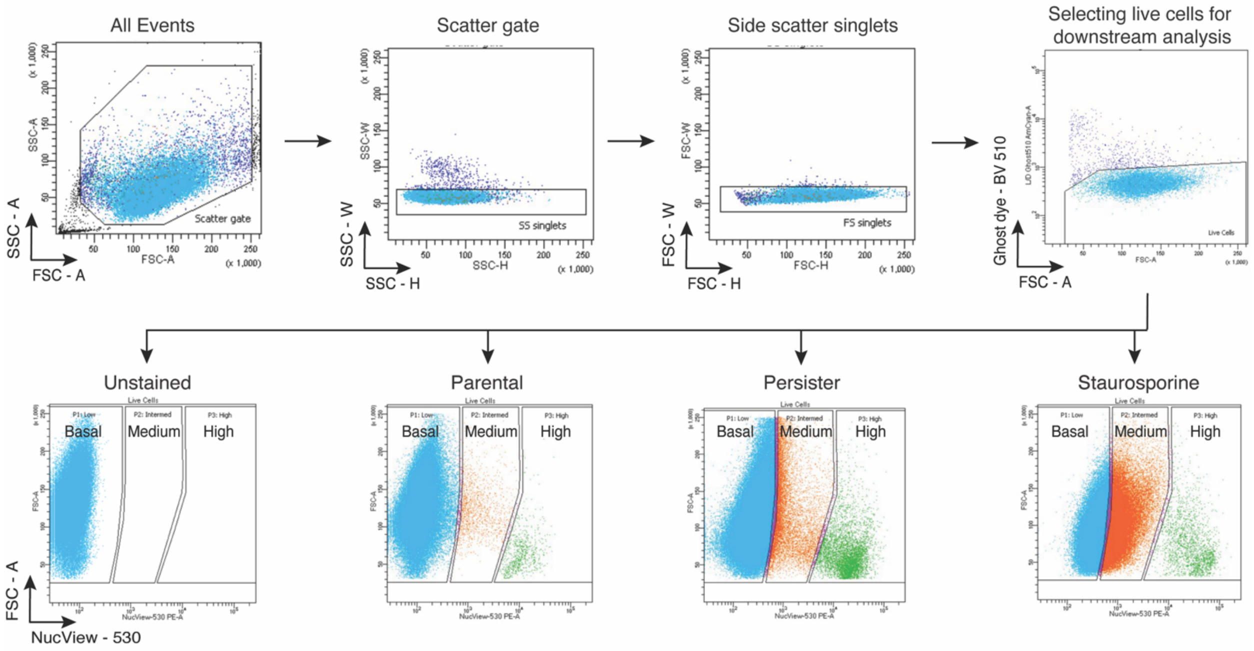


**Supplementary Figure 5: Caspase 3/7 activity reporter flow cytometry schematic. a**, Representative gating strategy for sorting caspase 3/7 activity sensor levels in A375 parental cells, persister cells treated with 250 nM dabrafenib and 25 nM trametinib, and staurosporine-treated positive control cells. Representative caspase 3/7 activity sensor levels for each condition shown in bottom row. This gating strategy was used to sort the Basal, Medium and High Caspase 3/7 activity level persister cell populations for Figure 1j,k and Extended Data Figure 2b,j.

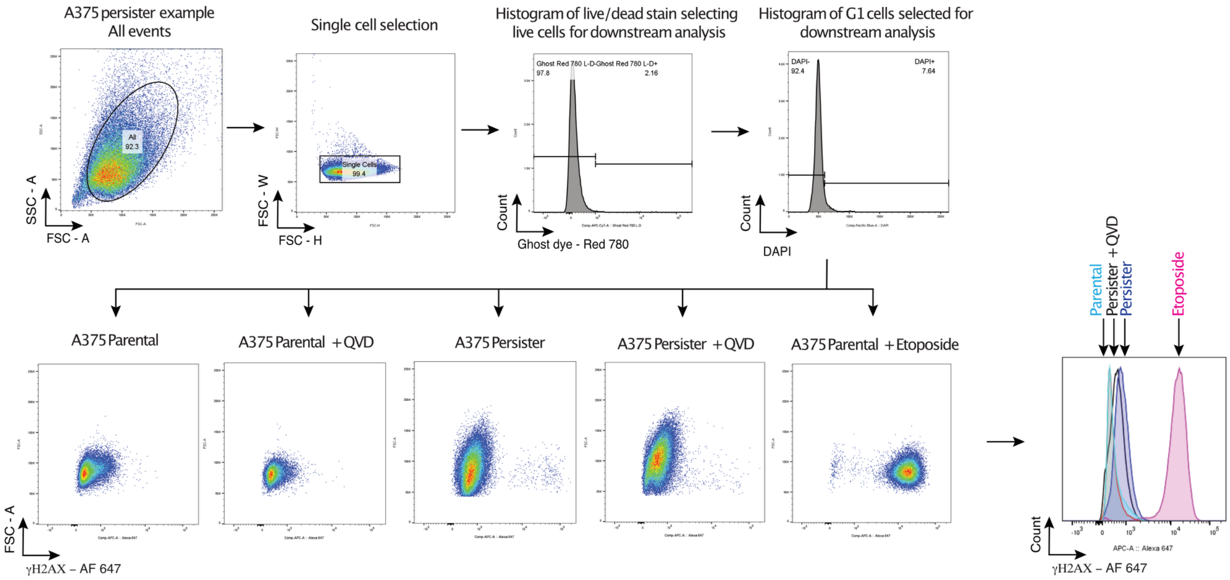


**Supplementary Figure 6: γH2AX DNA damage flow cytometry schematic.** Representative flow cytometry gating for γH2AX in A375 parental cells, persister cells treated with 250 nM dabrafenib and 25 nM trametinib, and positive control etoposide-treated cells. 10 µM QVD co-treatment was present for the duration of the experiment for QVD treated cells. Parental cell treatment with 100 µM etoposide for 30 minutes was used as a positive control. This gating was used for Extended Data Fig. 2h.

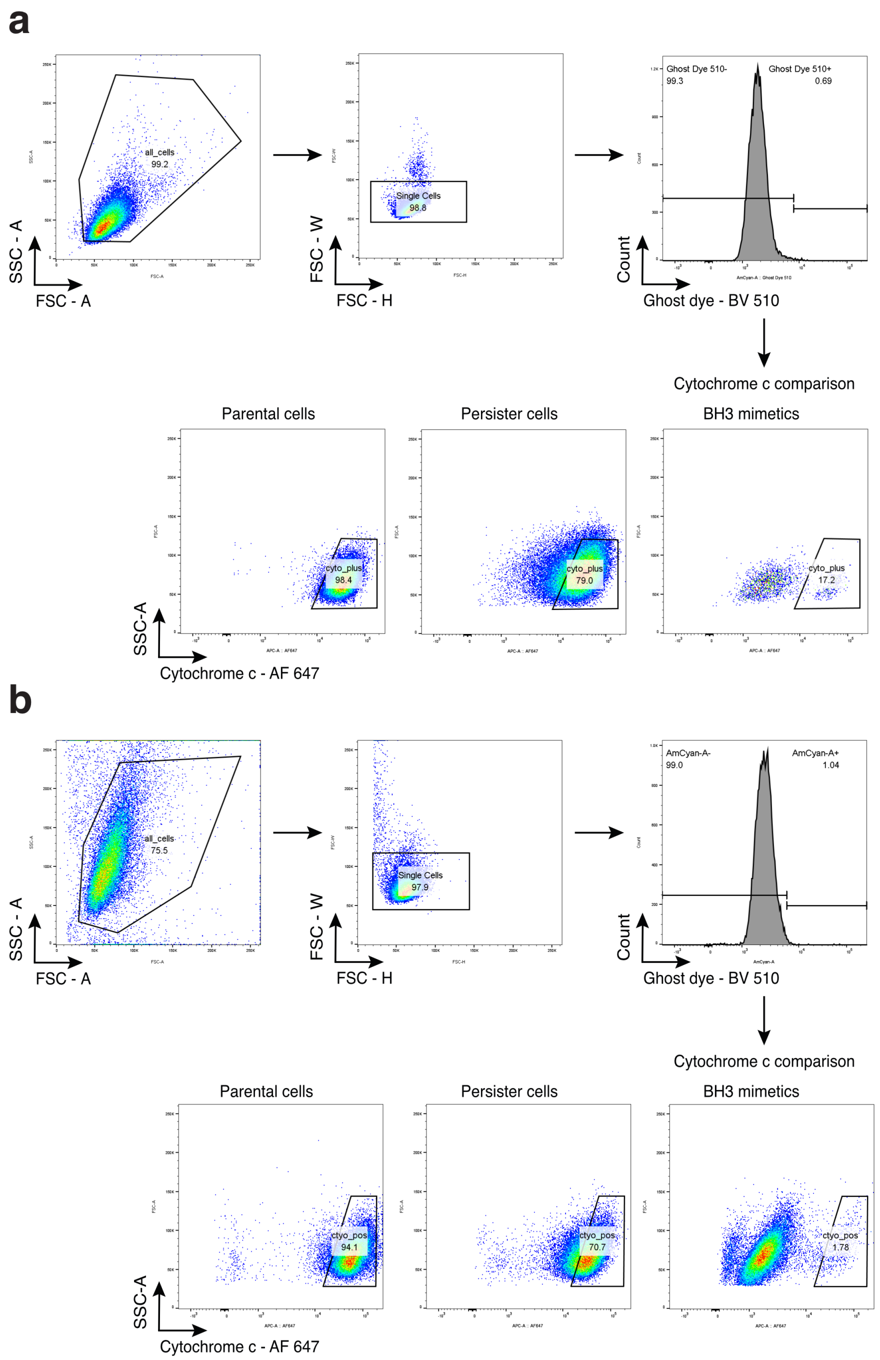


**Supplementary Figure 7: Mitochondrial release of cytochrome c flow cytometry assay with BH3 mimetics treatment schematic.** Gating strategy used for A375 (**a**) and PC9 (**b**) parental, BH3 mimetic-treated, and persister cells in Extended Data Fig. 7h-k. See methods for details of this assay.
